# A Molecular Odorant Transduction Model and the Complexity of Combinatorial Encoding in the *Drosophila* Antenna

**DOI:** 10.1101/237669

**Authors:** Aurel A. Lazar, Chung-Heng Yeh

**Affiliations:** Department of Electrical Engineering, Columbia University, New York, United States

## Abstract

In the past two decades, a substantial amount of work characterized the odorant receptors, neuroanatomy and odorant response properties of the early olfactory system of *Drosophila melanogaster.* Yet many odorant receptors remain only partially characterized and, the odorant transduction process and the axon hillock spiking mechanism of the olfactory sensory neurons (OSNs) have yet to be fully determined.

*Identity* and *concentration*, two key aspects of olfactory coding, originate in the odorant transduction process. Detailed molecular models of the odorant transduction process are, however, scarce for fruit flies. To address these challenges we advance a comprehensive model of fruit fly OSNs as a cascade consisting of an odorant transduction process (OTP) and a biophysical spike generator (BSG). We model identity and concentration in OTP using an odorant-receptor binding rate tensor, modulated by the odorant concentration profile, and an odorant-receptor dissociation rate tensor, and quantitatively describe the ligand binding/dissociation process. We model the BSG as a Connor-Stevens point neuron.

The resulting combinatorial encoding model of the *Drosophila* antenna provides a theoretical foundation for understanding the neural code of both odorant identity and odorant concentration and advances the state-of-the-art in a number of ways. First, it quantifies on the molecular level the combinatorial complexity of the transformation taking place in the antennae. The concentration-dependent combinatorial code determines the complexity of the input space driving olfactory processing in the downstream neuropils, such as odorant recognition and olfactory associative learning. Second, the model is biologically validated using multiple electrophysiology recordings. Third, the model demonstrates that the currently available data for odorant-receptor responses only enable the estimation of the affinity of the odorant-receptor pairs. The odorant-dissociation rate is only available for a few odorant-receptor pairs. Finally, our model calls for new experiments for massively identifying the odorant-receptor dissociation rates of relevance to flies.

## Introduction

The odorant response of olfactory sensory neurons (OSNs) in the *Drosophila* antennae has been experimentally characterized by multiple research groups (*de Bruyne et al., 2001*; *Hallem and Carlson, 2006*; *Smart et al., 2008*), and their results combined into a single consensus database, called the DoOR database (*Galizia et al., 2010*; *Münch and Galizia, 2016*). A key functionality of OSNs is to jointly encode both odorant identity and odorant concentration (*Masek and Heisenberg, 2008*; *Yarali et al., 2009*; *Nagel and Wilson, 2011*; *Wilson, 2013*; *Martelli et al., 2013*). A single odorant stimulus usually activates multiple OSN groups expressing the same receptor type, while different odorants activate different OSN groups (*Laurent, 1999*; *Hopfield, 1999*; *Touhara and Vosshall, 2009*). The identity of an odorant is combinatorially encoded by the set of responding OSN groups (*Malnic et al., 1999*), and the size of OSN set varies as the concentration changes (*Hallem and Carlson, 2006*). The temporal response of an OSN simultaneously represents the information of odorant concentration and concentration gradient also known as 2D odorant encoding (*Kim et al., 2011b, 2015*). These two aspects of olfactory coding, *identity* and *concentration*, originate in the odorant transduction process (*Hallem et al., 2004*; *Nagel and Wilson, 2011*). However, detailed molecular models of the odorant transduction process are scarce for fruit flies.

To address these challenges we advance a comprehensive model of fruit fly OSNs as a cascade consisting of an odorant transduction process (OTP) and a biophysical spike generator (BSG). We model identity and concentration in OTP by an odorant-receptor binding rate tensor, modulated by the odorant concentration profile, and an odorant-receptor dissociation rate tensor, and quantitatively describe the ligand binding/dissociation process. OSNs are distributed across the surface of maxillary palp and the third segment of antenna (de Bruyne et al., 2001; *Couto et al., 2005*; *Song et al., 2012*). Since there is no commonly accepted terminology in the literature for naming these two olfactory appendages as a single entity, and in order to avoid potential confusion, we will refer to the set of *all* OSNs on one side of the fly brain as an antenna/maxillary palp (AMP) local processing unit (LPU) (*Chiang et al., 2011*).

To biologically validate our modeling approach, we first propose an algorithm for estimating the affinity and the dissociation rate of an odorant-receptor pair. We then apply the algorithm to electrophysiology recordings and estimate the affinity and dissociation rate for three odorant-receptor pairs. Second, we evaluate the temporal response of the OSN model with a multitude of stimuli for all three odorant-receptor pairs. The output of the model closely reproduces the temporal responses of OSNs obtained from *in vivo* electrophysiology recordings (*Kim et al., 2011b, 2015*) for all three odorant-receptor pairs across all three types of stimuli.

Lastly, we evaluate the model at the OSN antennae population level. We first empirically estimate the odorant-receptor affinity using the spike count records in the DoOR database for 24 receptor types in response to 110 odorants (*Münch and Galizia, 2016*). With estimated affinity values, we simulate the temporal response of the OSN population to staircase odorant waveforms. The output of simulated OSN population demonstrates that the odorant identity is encoded in the set of odorant-activated OSN groups expressing the same receptor type, and, more importantly, the size of the set expands or reduces as the odorant concentration increases or decreases.

The fruit fly OSN model presented here provides a theoretical foundation for understanding the neural code of both odorant identity and odorant concentration. It advances the state-of-the-art in a number of ways. First, it models on the molecular level the combinatorial complexity of the transformation taking place in *Drosophila* antennae OSNs. The resulting *concentration-dependent combinatorial code* determines the complexity of the input space driving olfactory processing in the downstream neuropils, such as odorant recognition and olfactory associative learning. Second, the model is biologically validated using multiple electrophysiology recordings. Third, the OSN model demonstrates that the currently available data for odorant-receptor responses only enables the estimation of the affinity of the odorant-receptor pairs. Finally, our model calls for new experiments for massively identifying the odorant-receptor dissociation rates of relevance to flies.

## Model

Detailed biophysical models for the odorant transduction process have been proposed for worms and vertebrates. Such models are scarce for insects and, in particular, for fruit flies. Dougherty et al. proposed a frog odorant receptor model that exhibits a complex temporal response (*Dougherty et al., 2005*). Rospars et al. proposed a model that characterizes the steady state response of OSNs for rats and moths (*Rospars et al., 1996, 2008*). The model stands out for its simplicity and modeling clarity, while lacking temporal variability. Other notable models appeared in (Suzuki et al., 2002; *De Palo et al., 2012*). Recently, Cao et al. published a phenomenological model to characterize the peak and the steady response of sensory adaption for fruit fly OSNs (*Cao et al., 2016*). Gorur-Shandilya et al. proposed a two-state model for the fruit fly odorant receptors that can reproduce Weber-Fechner’s law observed in physiological recordings (*Gorur-Shandilya et al., 2017*). In addition, De Palo et al. (*De Palo et al., 2013*) proposed an abstract/phenomenological model with feedback mechanism that characterizes the common dynamical features in both visual and olfactory sensory transduction.

**Figure 1.**
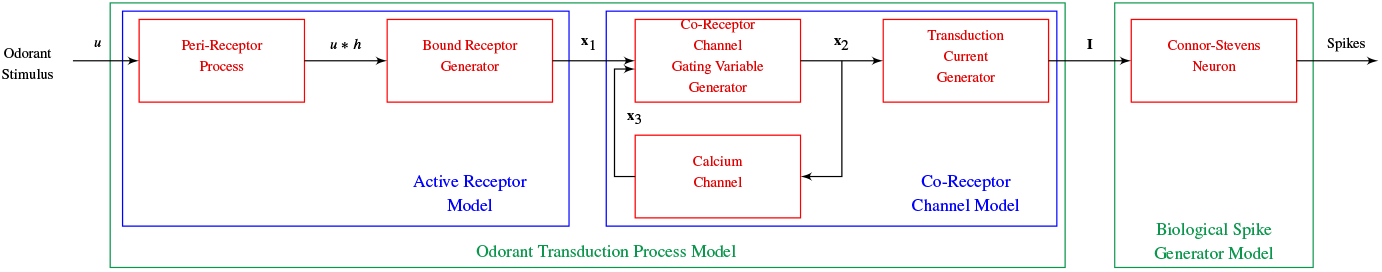
The block diagram of fruit fly OSN model consists of the OTP and the BSG.

Except for the transduction current recorded for studying sensory adaptation (*Cao et al., 2016*), reproducing the temporal response of the AMP LPU on either side of the brain has been scarcely investigated in the literature. In particular, 2D odorant encoding has not yet been successfully modeled. To address these challenges, we model the OSNs as a cascade consisting of an odorant transduction process (OTP) and a biophysical spike generator (BSG), as shown in Figure 1. The OTP model consists of an active receptor model and a co-receptor channel model (Larsson et al., 2004; *Benton et al., 2006*). The BSG model we employ here is based on the Connor-Stevens point neuron model (*Connor and Stevens, 1971*). The spike trains generated by the BSGs contain the odorant identity, odorant concentration, and concentration gradient information that the fly brain uses to make odorant valence decisions.

### The Odorant Transduction Process Model

Two research groups have published widely different results on the OR transduction process in fruit flies (Sato et al., 2008; *Wicher et al., 2008*; *Kaupp, 2010*). As the exact signaling of the transduction cascade in fruit flies is still inconclusive, our approach focusses here on constructing a minimal transduction model. Called the fruit fly odorant transduction process (OTP) model, it extends the model proposed by (*Rospars et al., 1996, 2008*) by incorporating the essential features of temporal dynamics of other computational models, such as the one proposed by (*Dougherty et al., 2005*), while at the same time exhibiting the calcium dynamics of (*Cao et al., 2016*). In the latter work, the temporal dynamics of fly’s OSN vanish in the absence of extracellular calcium. Notably, the calcium dynamics considered here constitutes a feedback mechanism that is similar to but also different from the one in the abstract model proposed by (*De Palo et al., 2013*).

Olfactory transduction in fruit flies from airborne molecules to transduction current involves a number of steps (*Kaissling, 2013*; *Leal, 2013):* i) absorption of odorant molecules through the sensillum surface, binding between odorant molecules and odorant-binding proteins (OBPs), and diffusion of bound OBPs through the aqueous sensillar lymph to OSN dendrites, ii) odorant-receptor binding/dissociation, and iii) opening of ion channels that results in transduction current. The first step is known as the “peri-receptor” processing, the second step is referred as the bound receptor generator and the third step as the co-receptor channel. Taken together, they represent the fruit fly odorant transduction process.

**Figure 2.**
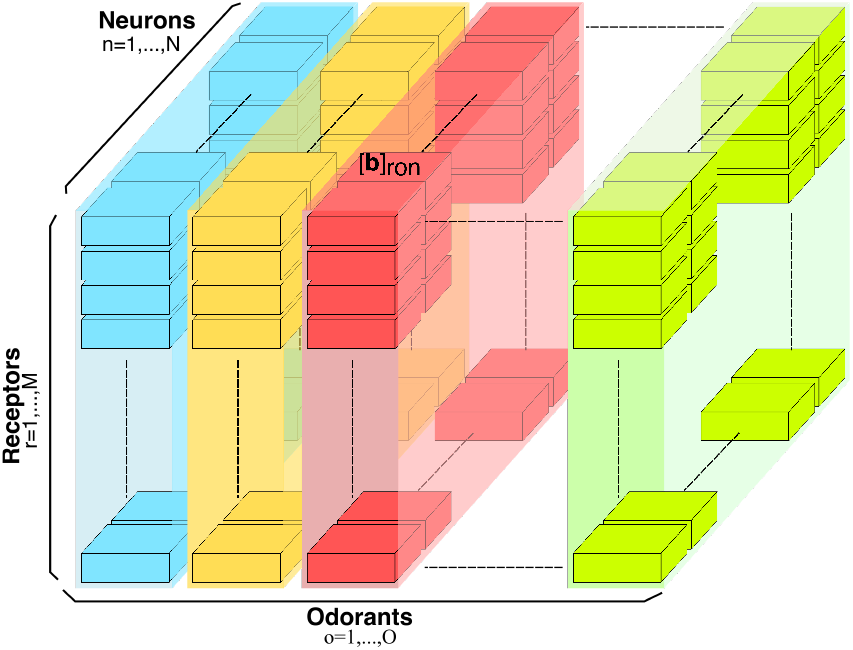
Three dimensional odorant-receptor binding rate tensor **b**. For a given neuron *n* = 1,2, &,*N*, the binding rate values are denoted by [**b**]_*ron*_, for all *r* = 1,2,&, *R*, and *o* = 1,2,&, *O*. For the fruit fly, the total number of neurons expressing the same receptor type is about *N* = 25, and the total number of receptor types is around *R* = 60. *O* is the number of all odorants that the fruit fly senses.

We propose an olfactory transduction process model that consists of an *active receptor* model and a *co-receptor channel* model (*Larsson et al., 2004*; *Benton et al., 2006*). The active receptor contains a peri-receptor model and a bound-receptor model. The peri-receptor process is modeled as a linear filter that describes the transformation of an odorant concentration waveform as odorant molecules diffuse through sensilla walls towards the OSN dendrites. The bound-receptor model encodes odorant identity and odorant concentration with a binding rate tensor, modulated by the odorant concentration profile, and a dissociation rate tensor. The odorant concentration profile is defined as the linearly weighted sum of the filtered odorant concentration and the filtered concentration gradient. Modulation is modeled here as a product. The co-receptor channel represents the ion channel gated by the atypical co-receptor (CR), Or83b. The calcium channel models the calcium dynamics, and provides a feedback mechanism to the co-receptor channel. The transduction current generator models the transmembrane current through the co-receptor channel.

#### The Active Receptor Model

The fruit fly active receptor model quantifies the binding and the dissociation process between odorant molecules and odorant receptors. As introduced here, the model centers on the rate of change of the ratio of free receptors versus the total number of receptors [**x**_0_]_*ron*_ expressed by neuron *n*:

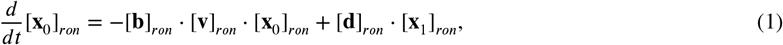

where [**x**_1_]_*ron*_ is the ratio of ligand-bound receptors. Here *r* = 1,2,*R*, is the receptor type, *o* = 1,2&, *O*, denotes the odorant and *n* = 1,2,&, *N*, denotes the neuron index. In Equation 1 above the ratios [**x**_0_]_*ron*_ and [**x**_1_]_*ron*_ are entries of the 3D tensors **x**_0_ and **x**_1_, respectively. The 3D tensor **b** with entries [**b**]_*ron*_ is called the *odorant-receptor binding rate* and models the association rate between an odorant and a receptor type. The 3D tensor **d** with entries [**d**]_*ron*_ denotes the odorant-receptor dissociation and models the detachment rate between an odorant and a receptor type. The 3D binding rate tensor **b** is graphically depicted in Figure 2 (A similar figure can be drawn for the dissociation rate tensor **d**). In what follows, the biding rate [**b**]_*ron*_ and the dissociation rate [**d**]_*ron*_, for a given odorant *o* and a given receptor type *r*, are assumed for simplicity to take the same value for all neurons *n* = 1, 2,&, *N*.

**Table 1.** Summary of the variables in the fruit fly odorant transduction model.

| Variable | Description |
| --- | --- |
| $u$ | odorant concentration waveform |
| $v$ | odorant concentration profile |
| $\mathbf{b}$ | odorant-receptor binding rate |
| $\mathbf{d}$ | odorant-receptor dissociation rate |
| $\mathbf{x}_1$ | ratio of ligand-bound receptors |
| $\mathbf{x}_2$ | gating variable of the *co-receptor* channel |
| $\mathbf{x}_3$ | state variable of the calcium channel |
| $\mathbf{I}$ | transduction current generated by the *co-receptor* channel |

The tensor **v** with entry [**v**]_*ron*_ in Equation 1 is the odorant concentration profile and it is given by

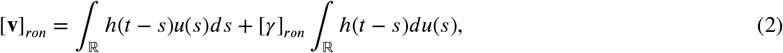

if the RHS is positive and zero otherwise. The RHS is the weighted sum of the filtered odorant concentration *u* and the filtered concentration gradient *du/dt* with [*γ*]_*ron*_ denoting a weighting factor. The impulse response of the linear filter *h* = *h*(*t*), 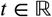 (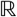 denotes the set of real numbers), models the “peri-receptor” process that describes the transformation of odorant concentration waveform as odorant molecules diffuse through sensilla walls towards the dendrites of OSN (*Fleischer et al., 2017*). For simplicity, the dependence of *h*(*t*) on the geometry of the sensillum and the diffusion of odorant molecules across the sensillar lymph is not considered here. *h*(*t*) in Equation 2 is the impulse response of a low-pass linear filter that is usually defined in the literature in frequency domain. Alternatively, *h*(*t*) is the solution to the second-order differential equation, see **Appendix 1**.

Note that the odorant transduction models in the literature only consider the odorant concentration but not the odorant concentration profile as the input to the transduction cascade (*De Palo et al., 2012, 2013*; *Rospars et al., 1996, 2008*; *Cao et al., 2016*). As we will show, the odorant concentration profile is critical for modeling 2D odorant encoding.

We assume that receptors only have two states, either being “free” or “bound”, *i.e.*, [**x**_0_]_*ron*_ + [**x**_1_]_*ron*_ = 1. Then, Equation 1 amounts to,

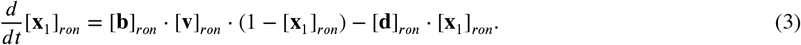

Equation 3 maps the input given by the product between the binding rate and the odorant concentration profile, and the dissociation rate, i.e., ([**b**]_*ron*_ · [**v**]_*ron*_, [**d**]_*ron*_), into the ratio of bound receptors [**x**_1_]. In what follows this map will be called the *bound-receptor generator.*

#### The Co-Receptor Channel Model

The fruit fly co-receptor channel and a calcium channel appear in a feedback configuration. Each of these components has its specific functionality. The *co-receptor channel* represents the ion channel gated by the atypical co-receptor (CR), Or83b. The *calcium channel* models the calcium dynamics, and provides a feedback mechanism to the co-receptor channel.

Next, we walk through each of the three equations of the co-receptor channel model. The key variables involved in the proposed odor transduction model are summarized in Table 1.

1. The rate of change of *the gating variable of the co-receptor channel* [**x**_2_]_*ron*_:

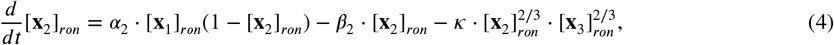

where *α*_2_ and *β*_2_ are scalars indicating the rate of activation and deactivation of the gating variable, respectively, and 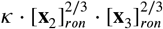 models the calcium feedback with *κ* a constant. The co-receptor channel model considered here differs from the one proposed by De Palo et al. (*De Palo et al., 2013*) in two important ways. First, the input to the co-receptor channel is the ratio of the ligand-bound receptors [**x**_1_]_*ron*_ driven, among others, by the odorant concentration profile [**v**]_*ron*_, while De Palo et al. used the odorant concentration *u*. Second, the feedback mechanism is based on the fractional power 2/3 for the variables [**x**_2_]_*ron*_ and [**x**_3_]_*ron*_, while De Palo et al. used the variables raised to power 1 in their feedback model. The fractional power is key in facilitating the encoding of the filtered concentration gradient.
2. The rate of change of *the state* variable of the calcium channel [**x**_3_]_*ron*_:

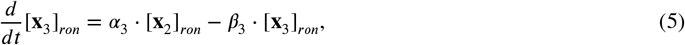

where *α*_3_ and *β*_3_ are scalars indicating the rate of increase and decrease of the state variable.
3. Finally, the transduction current [**I**]_*ron*_ is given by:

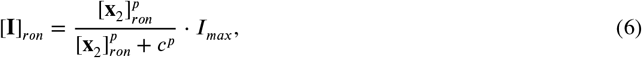

where *p* and *c* are scalars, and *I_max_* denotes the maximal amplitude of the current through the *co-receptor* channel, whose value is empirically determined through parameter sweeping. Combining the equations introduced above, we rewrite the odorant transduction process model in compact form as

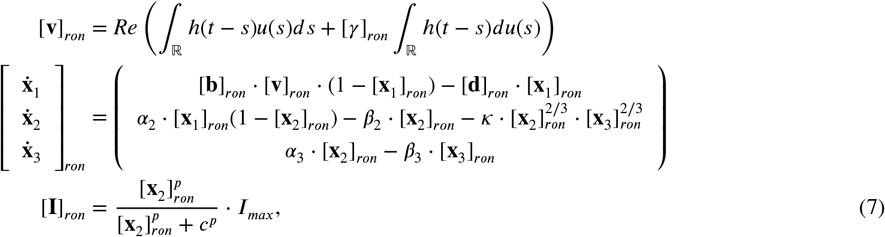

for all *r* =1, 2,&, *R, o* = 1, 2,&, *O* and *n* = 1,2,&, *N*. *Re* above denotes the rectification function.

### Biophysical Spike Generator Model

We restrict our choice of the spiking mechanism of OSNs to biophysical spike generators (BSG) such as the Hodgkin-Huxley, the Morris-Lecar, and the Connor-Stevens point neuron models. For simplicity of presentation, we only consider in Appendix 2 the Connor-Stevens (CS) point neuron model.

## Results

### Biological Validation of the OSN Model

The essential functionality of OSNs is to jointly encode both odorant identity and odorant concentration. To address these two functional aspects we modeled each OSN as an OTP/BSG cascade. To validate our approach, we examine here the response of the OSN model to odorant waveforms that were previously used in experiments with different odorants and receptors, and compare the model responses with electrophysiological recordings.

We first estimate the affinity and dissociation rate for the (*acetone, Or59b*) pair using two different datasets of electrophysiology recordings. Second, we evaluate the temporal response of the *Or59b* OSN model to acetone with a multitude of stimuli, including step, ramp and parabola waveforms. We further interrogate the model with staircase and white noise waveforms. Our results show that the model closely matches the complex temporal response of *Or59b* OSNs from electrophysiological recordings. Lastly, we evaluate the affinity and dissociation rate for different odorant-receptor pairs including (*methyl butyrate, Or59b*) and (*butyraldehyde, Or7a*).

**Figure 3.**
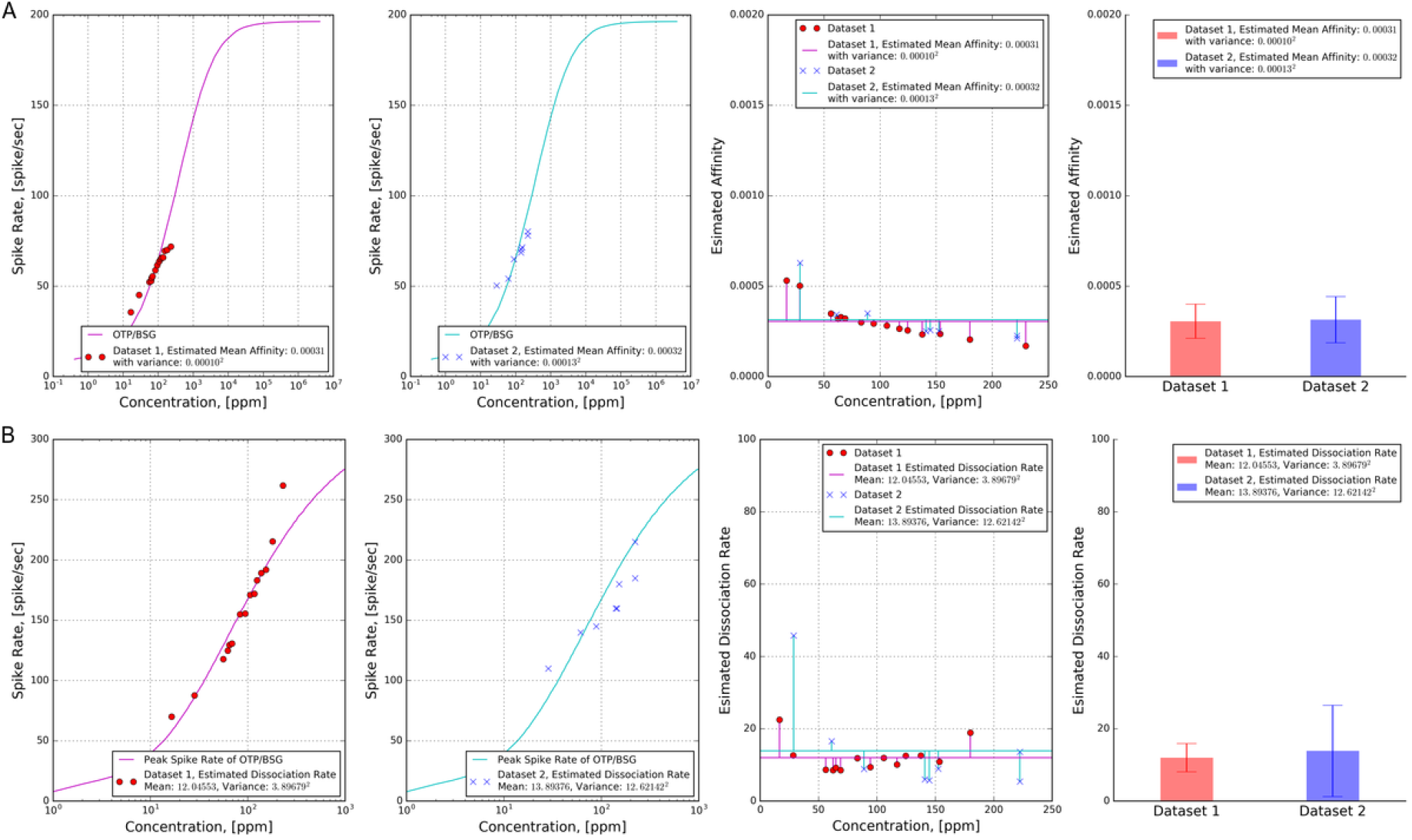
(**A**) Estimation of the affinity value for two datasets. Both datasets contain PSTHs of OSNs expressing *Or59b* in response to acetone step waveforms. The source of the two datasets is given in **Appendix 3**. For each of the datasets, we computed the mean and variance of the affinity empirically estimated for each data point. (**Left 1**) Dataset 1: Estimated affinity is 3.141 · 10^−4^ with variance (1.312 · 10^−4^)^2^; (**Left 2**) Dataset 2: Estimated affinity is 3.201 · 10^−4^ with variance (1.001 · 10^−4^)^2^; (**Right 2**) Estimation of the affinity as a function of concentration amplitude. (**Right 1**) The mean and variance of estimated affinity value. (**B**) Estimation of the dissociation rate for two datasets. Both datasets contain PSTHs of OSNs expressing *Or59b* in response to acetone step waveforms. The source of the two datasets is given in **Appendix 3**. For each of the datasets, we computed the mean and variance of the dissociation rates empirically estimated for each data point. (**Left 1**) Dataset 1: Estimated dissociation rate is 1.205 · 10^1^ with variance (3.900 · 10^1^)^2^; (**Left 2**) Dataset 2: Estimated dissociation rate is 1.389 · 10^1^ with variance (1.262 · 10^1^)^2^; (**Right 2**) Estimation of the dissociation rate as a function of concentration amplitude. (**Right 1**) The mean and variance of estimated dissociation rate.

#### Estimating the Affinity, Binding and Dissociation Rates for (*Acetone, Or59b*)

We applied **Algorithm 1** (see *Methods and Materials* section) to estimate the affinity, the dissociation rate, and the binding rate for (*acetone, Or59b*) by using two different datasets of electrophysiology recordings. The source of the two datasets is given in **Appendix 3**. Each of the two datasets contains the PSTHs obtained from the response of OSNs expressing *Or59b* to acetone step waveforms with different concentration amplitudes. As required by **Algorithm 1**, we first retrieved the peak and steady state spike rates from the PSTH in response to each concentration amplitude recorded in the datasets. Second, for each of the two datasets, we used the steady state spike rate to estimate the value of the affinity for each concentration amplitude, and computed the mean and variance of the affinity as shown in Figure 3.A. With the mean of the estimated affinity, we then used the peak spike rate to estimate the value of the dissociation rate for each concentration amplitude, and computed the mean and variance of the dissociation rate as shown in Figure 3.B. For the first dataset, the mean and variance of the estimated affinity are 3.141 · 10^−4^ and (1.312 · 10^−4^)^2^, respectively, and the mean and variance of the estimated dissociation rate are 1.205 · 10^1^ and (3.900 · 10^1^)^2^, respectively. For the second dataset, the mean and variance of the estimated affinity are 3. 201 · 10^−4^ and (1.001 · 10^−4^)^2^, respectively, and the mean and variance of the estimated dissociation rate are 1.389 · 10^1^ and (1.262 · 10^1^)^2^, respectively. The values of the affinity estimated from the two datasets are almost identical, while the two estimated dissociation rates are marginally different. This is because the steady state spike rates of the two datasets are similar, but the peak spike rates of the two datasets differ slightly, as shown in Figure 1 in **Appendix 3**.

#### Evaluating the Temporal Response of the *Or59b* OSN Model to Acetone

To evaluate temporal response, we stimulated the OSN model with multiple odorant stimuli that were previously used in experiments designed for characterizing the response to *acetone* of OSNs expressing *Or59b.* For all OTP models considered below we set the odorant-receptor binding rate to 3.141 · 10^−4^ and the odorant-receptor dissociation rate to 1.205 · 10^1^.

#### Response of the *Or59b* OSN Model to Step, Ramp and Parabola Acetone Waveforms

**Figure 4.**
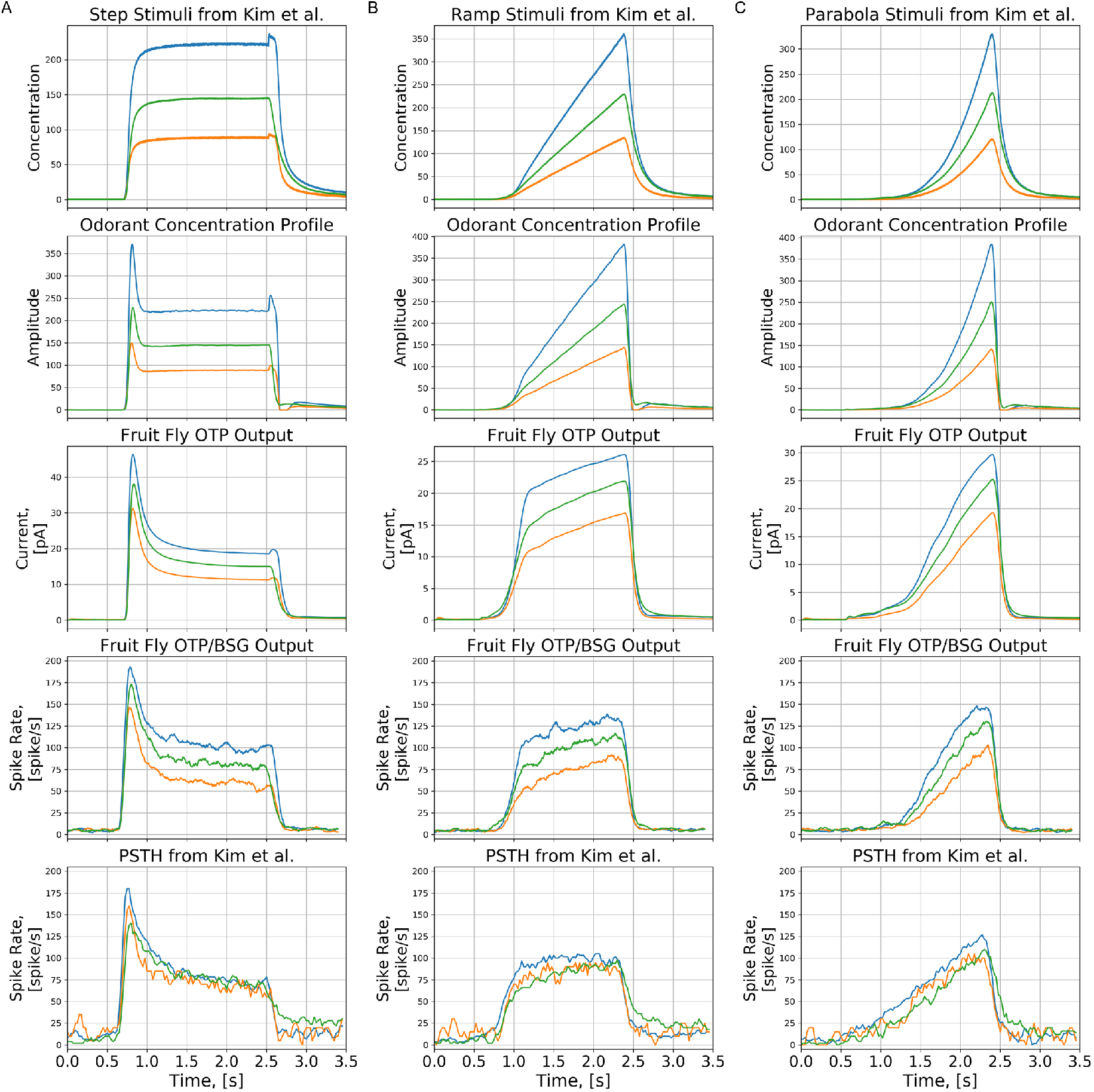
Characterization of the OTP/BSG cascade in response to step, ramp, and parabola stimuli. Odorant: *acetone*, receptor: *Or59b.* The stimulus waveforms are identical to the ones used in (*Kim et al., 2015*). The odorant-receptor binding and dissociation rates were set to 3.141 · 10^−4^ and 1.205 · 10^1^. (**A**) Step stimuli. (**B**) Ramp stimuli. (**C**) Parabola stimuli. (**First row**) Stimulus waveforms. (**Second row**) The odorant concentration profile. (**Third row**) The transduction current at the output of the OTP model. (**Forth row**) PSTH computed from the output of the OTP/BSG output. (**Fifth row**) PSTH of the spike train generated by the *Or59b* OSN in response to the stimulus waveforms (Reproduced from Figure 2 in (*Kim et al., 2015*) using the original raw data).

We first evaluated the response of the *Or59b* OSN model to step, ramp, and parabola stimulus waveforms as shown in the first row of Figure 4. The temporal response of the OTP/BSG cascade (the fourth row of Figure 4) is similar to the one of the OTP model (the third row of Figure 4). For step stimuli, the OTP/BSG cascade generates a chair-shaped response by first picking up the gradient of the concentration right after the onset of the odorant, and then gradually dropping down to a constant value, that encodes the step value of the amplitude. For ramp stimuli, the initial response of the OTP/BSG cascade rapidly increases, and then it plateaus as the gradient of ramp stimuli becomes constant. Lastly, for the parabola stimuli, the response of the OTP/BSG cascades resembles a ramp function, that corresponds to the gradient of parabola stimulus waveforms.

Furthermore, we also compared the PSTH of the spike trains generated by the OTP/BSG cascade with the PSTH of an *Or59b* OSN obtained from electrophysiology recordings in response to acetone concentration waveforms (*Kim et al., 2015*). As shown in Figure 4, the OTP/BSG closely matches the odorant response of the *Or59b* OSN.

#### Response of the *Or59b* OSN Model to White Noise *Acetone* Waveforms

To further compare the response of the OTP/BSG cascade with the *Or59b* OSN response, we stimulated OTP/BSG cascades with white noise stimuli, and compared the PSTH of the model with the one from experimental recordings. The white noise stimulus was previous used in the experimental setting of (*Kim et al., 2011b*) for characterizing the response of *Or59b* OSNs to acetone.

**Figure 5.**
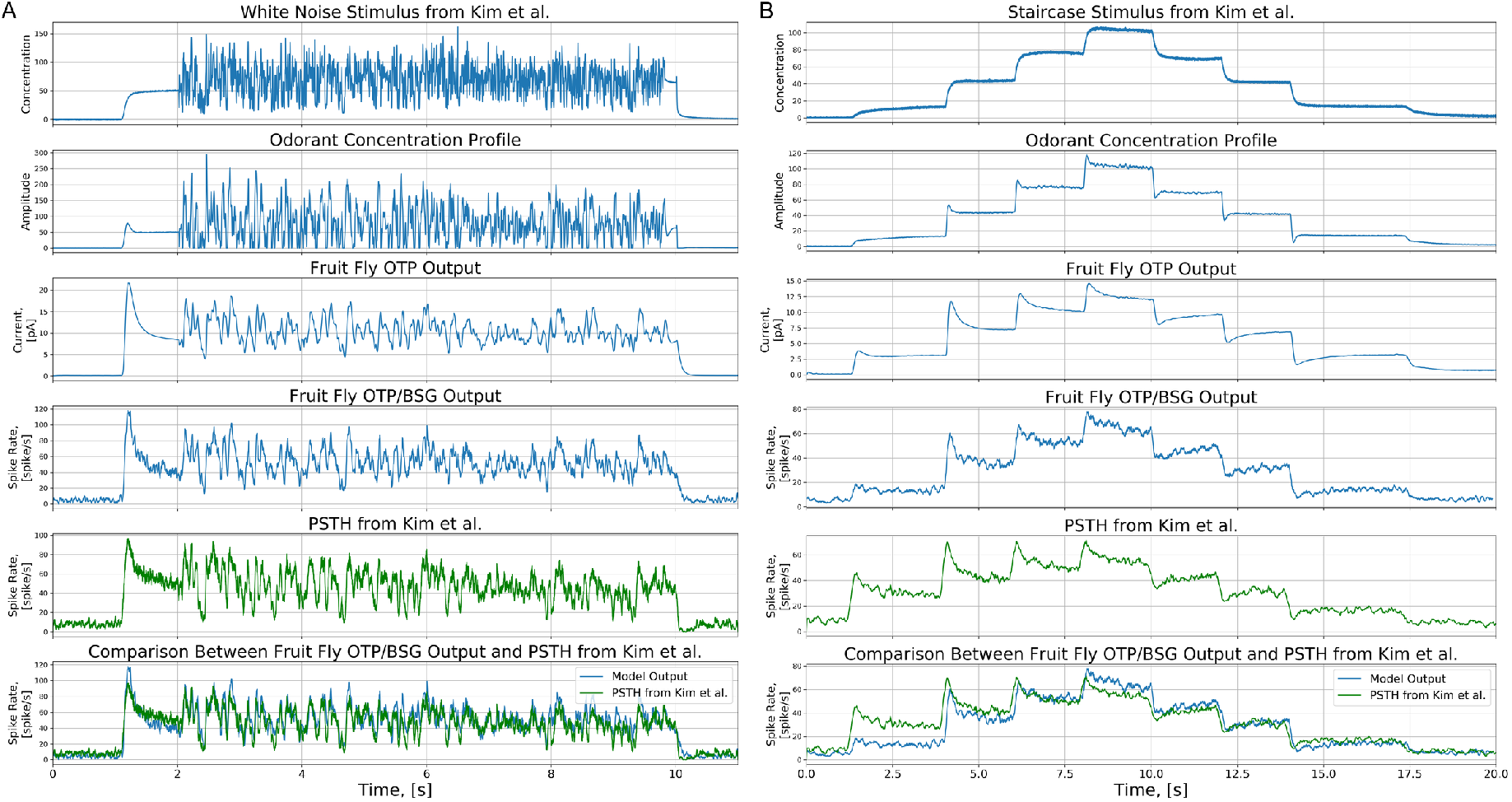
Characterization of the OTP/BSG cascade in response to white noise and staircase stimuli previously used in (*Kim et al., 2011b*). Odorant: *acetone*, receptor: *Or59b.* The odorant-receptor binding and dissociation rates were set to 3.141 · 10^−4^ and 1.205 · 10^1^. (**A**) White noise. (**B**) Staircase. (**First row**) White noise and staircase stimuli. (**Second row**) The odorant concentration profile. (**Third row**) The output of the OTP model. (**Forth row**) The PSTH of the spike train generated by the OTP/BSG cascade. (**Fifth row**) The PSTH of the spike train of the recorded OSN (Reproduced from Figure 2 (staircase) and 4 (white noise) in (*Kim et al., 2011b*) using the original raw data). (**Sixth row**) Comparison between the PSTHs at the output of the OTP/ BSG cascade and the recorded OSN.

The output of each of the stages of the *Or59b* OSN model are shown in Figure 5.A. The odorant onset at around 1 second is picked up by the odorant concentration profile (see the the second row of Figure 5.A. In addition, the white noise waveform between 2 and 10 second is smoothed out. The smoothing effect is due to the peri-receptor process filter. The OTP model further emphasizes the gradient encoding (the third row of Figure 5.A), and predominantly defines the temporal response of the OSN model to white noise stimuli. The BSG output follows the OTP output, as the BSG is simply a sampling device. Lastly, we compare the model output and the PSTH from OSN recordings in (*Kim et al., 2011b*) (the fifth and the sixth rows of Figure 5.A). The *Or59b* OSN model output PSTH closely matches the PSTH obtained from recordings.

The peri-receptor process filter is critical in processing the white noise waveforms, but less critical in processing the static waveforms discussed in this work. The filter prevents the model from overemphasizing the gradient of the white noise waveforms. In absence of this filter, the response of the OTP/BSG cascade is severely limited in matching the response of *Or59b* OSN (*Kim et al., 2011b*) to acetone waveforms.

#### Response of the *Or59b* OSN Model to Staircase *Acetone* Waveforms

Next, we stimulated OTP/BSG cascade with the staircase waveform that was previously used in experiments (*Kim et al., 2011b*), evaluated the PSTH from the resultant spike sequences, and compared the model PSTH to the one from experimental recordings.

As shown in the second row of Figure 5.B, the filter h(t) in Equation 2 has negligible effect on the odorant concentration profile since the staircase is smooth unlike the white noise stimulus discussed above. The encoding at jump times is strongly sharpened by the OTP. Overall, the fruit fly OTP/BSG cascade indeed encodes both the concentration and concentration gradient. In particular, at each upward concentration jump, the PSTH of the OSN launches upon a local maximum and then drops down to a saturation point. In addition, at each downward concentration jump, the same PSTH drops down first to a local minimum and then bounces back.

In short, the OSN model closely reproduces the temporal response of *Or59b* OSNs for all tested stimuli. This suggests that the OPT/BSG cascade has the desired complexity to effectively model the fruit fly OSNs.

**Figure 6.**
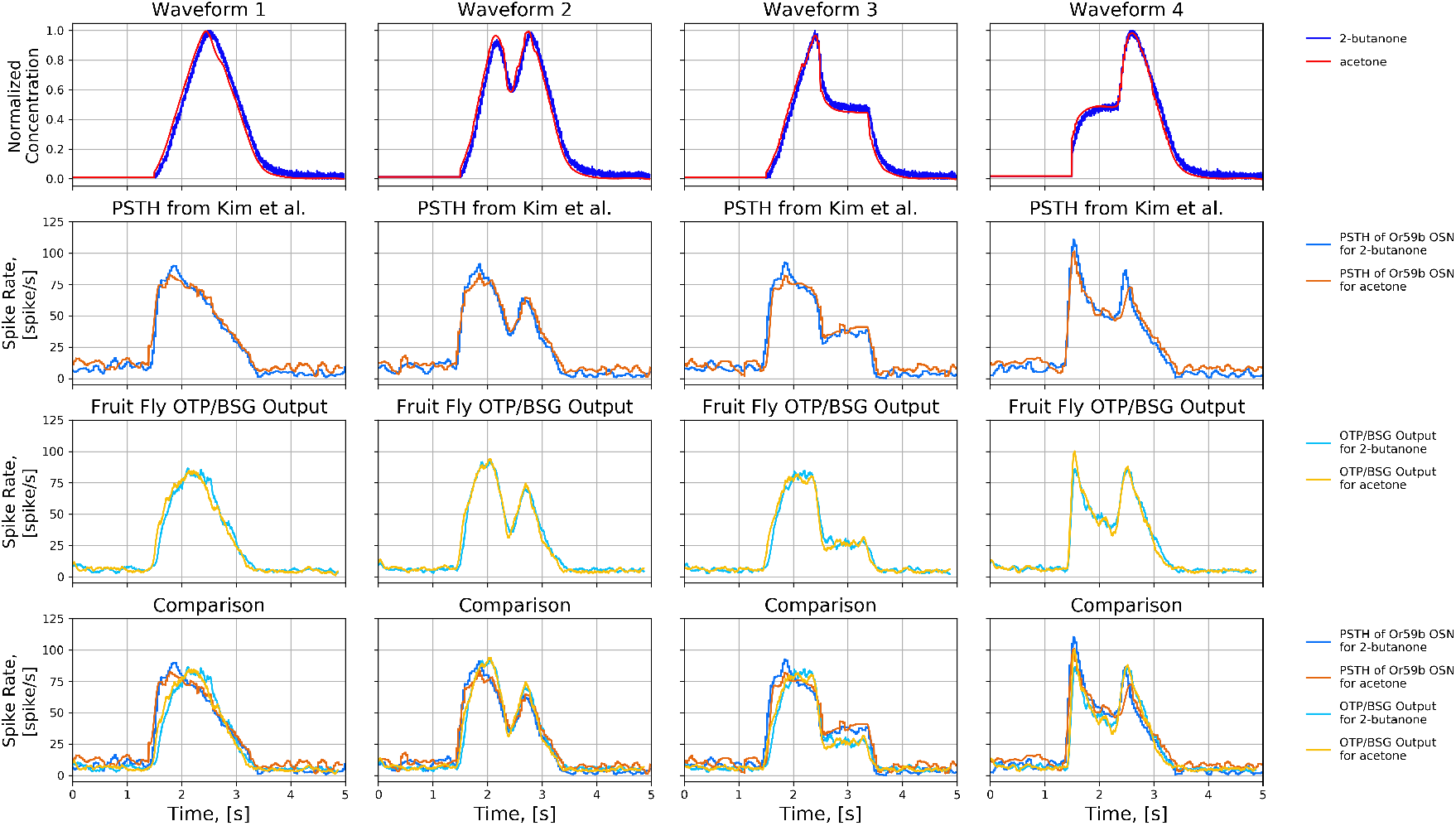
Comparison of the responses of an Or59b OSN to four concentration waveforms of *acetone* and *2-butanone.* (**First row**) Four normalized concentration waveforms. Each normalized waveform is scaled by 100 for *acetone* and by 10 for *2-butanone.* (**Second row**) PSTH of the *Or59b* OSN in response to the two odorants (Reproduced from (*Kim et al., 2011a*) using the original raw data.) (**Third row**) The PSTH of the spike train generated by the OTP/BSG cascade. (**Forth row**) Comparison between the PSTHs at the output of the OTP/BSG cascade and the recorded OSN in (*Kim et al., 2011a*).

#### Evaluating Affinity, Binding and Dissociation Rates of Other (*Odorant, Receptor*) Pairs

We next interrogate the role of the binding and dissociation rates in the OTP/BSG cascade. For a given receptor type and two odorants with different binding rates and the same dissociation rate, responses of the OTP/BSG cascade are identical if the waveforms of two odorants only differ by a scaling factor that is the reciprocal of the ratio of the biding rates. This follows from Equation 3.

#### Response of *Or59b* OSNs to Two Different Odorants

We verify the prediction mentioned above by stimulating the *Or59b* OSN model with two odorant stimuli, *acetone* and *2-butanone*, paired with four concentration waveforms, and compare the responses with the experimental recordings in (*Kim et al., 2011a*). As shown in the first row of Figure 6, the two odorants have identical normalized waveforms scaled by two different factors, 100 and 10. The affinity of *acetone* and *2-butanone* were estimated to be 3.141 · 10^−4^ and 3.164 · 10^−3^, respectively, and the dissociation rate of the two odorants were estimated to be 1.205 · 10^1^ and 1.203 · 10^1^, respectively.

The two odorant stimuli elicit almost exactly the same response from the *Or59b* OSN recodings (see the second row of Figure 6) as well as from the output of the OTP/BSG cascade (see the third row of Figure 6). The difference in binding rate for *acetone and2-butanone* is perfectly counterbalanced by the scaling factors of the odorant waveforms in Figure 6. In addition, the output of the OTP/BSG cascade closely reproduces the PSTH of the *Or59b* OSN as shown in the forth row of Figure 6.

#### Evaluating the Odorant-Receptor Response of the OTP/BSG Cascade

We further investigated the role of the binding rate using three odorant-receptor pairs that were previously used in experimental settings (*Kim et al., 2011a*). In addition to the binding rate estimated for (*acetone*, *Or59b*), we applied **Algorithm 1** to two additional odorant-receptor pairs using the original raw data presented in (*Kim et al., 2011a*):

1. for (*methyl butyrate, Or59b*) an affinity value 4.264 · 10^−4^ and a dissociation value 3.788 · 10^0^ were obtained from the steady-state spike rate at 87 spikes per second and the peak spike rate at 197 spikes per second in response to a constant stimulus with amplitude 20 ppm;
2. for (*butyraldehyde, Or7a*) an affinity value of 7.649 · 10^−3^ and a dissociation value 8.509 · 10^0^ were obtained from the steady-state spike rate at 43 spikes per second and the peak spike rate at 101 spikes per second in response to a constant stimulus with amplitude 173 ppm;

We simulated the OSN model for each of the three odorant-receptor pairs with three types of stimuli, step, ramp, and parabola. The binding and dissociation rates for different odorant-receptor pairs above were separately set.

As shown in Figure 7, with only the change in the value of the binding and dissociation rates, the OSN model closely matches the OSN’s response for all three tested odorant-receptor pairs. The results in Figure 7 suggests that a pair of binding and dissociation rates is capable of closely matching the temporal response of different temporal odorant concentration waveforms.

### Estimating the *Odorant-Receptor* Affinity Matrix with DoOR Datasets

The DoOR database integrates OSN recordings obtained with different measurement techniques (*Galizia et al., 2010*; *Münch and Galizia, 2016*), including *in situ* spike counts (*De Bruyne et al., 1999*; *Hallem et al., 2004*; *Smart et al., 2008*) and calcium activity (*Pelz et al., 2006*), among others. *Spike counts* are directly available from OSN spike train recordings. Relating calcium activity to spike activity is, however, error prone. We consequently focus here on the odorant-OSN response datasets of the DoOR database that contain spike count information (*Hallem and Carlson, 2006*). These datasets currently contain spike counts of 24 OSN groups in response to 110 odorants with a constant amplitude of 100 pm. The spike count is color coded and depicted Figure 8.A. By employing **Algorithm 1**, we empirically estimated the affinity value for all 110 · 24 = 2, 640 odorant-receptors pairs. The estimated affinity corresponding to each entry of the spike rate matrix shown in Figure 8.A is depicted in Figure 8.B.

In summary, the binding and dissociation rate model together with the rest of the OTP/BSG cascade define a family of OSN models, and provide the scaffolding for studying the neural coding for odorant *identity* and odorant *concentration* in temporal domain at the OSN antennae population level.

### Evaluating the Temporal Response of the AMP LPU

We investigated the temporal response of the OTP/BSG cascade to various odorant waveforms, including step, ramp, parabola, staircase, and white noise waveforms. In addition, we biologically validated the cascade with electrophysiological recordings of OSNs by demonstrating that the cascade is capable of reproducing the complex temporal responses of OSNs for multiple odorant receptor pairs.

**Figure 7.**
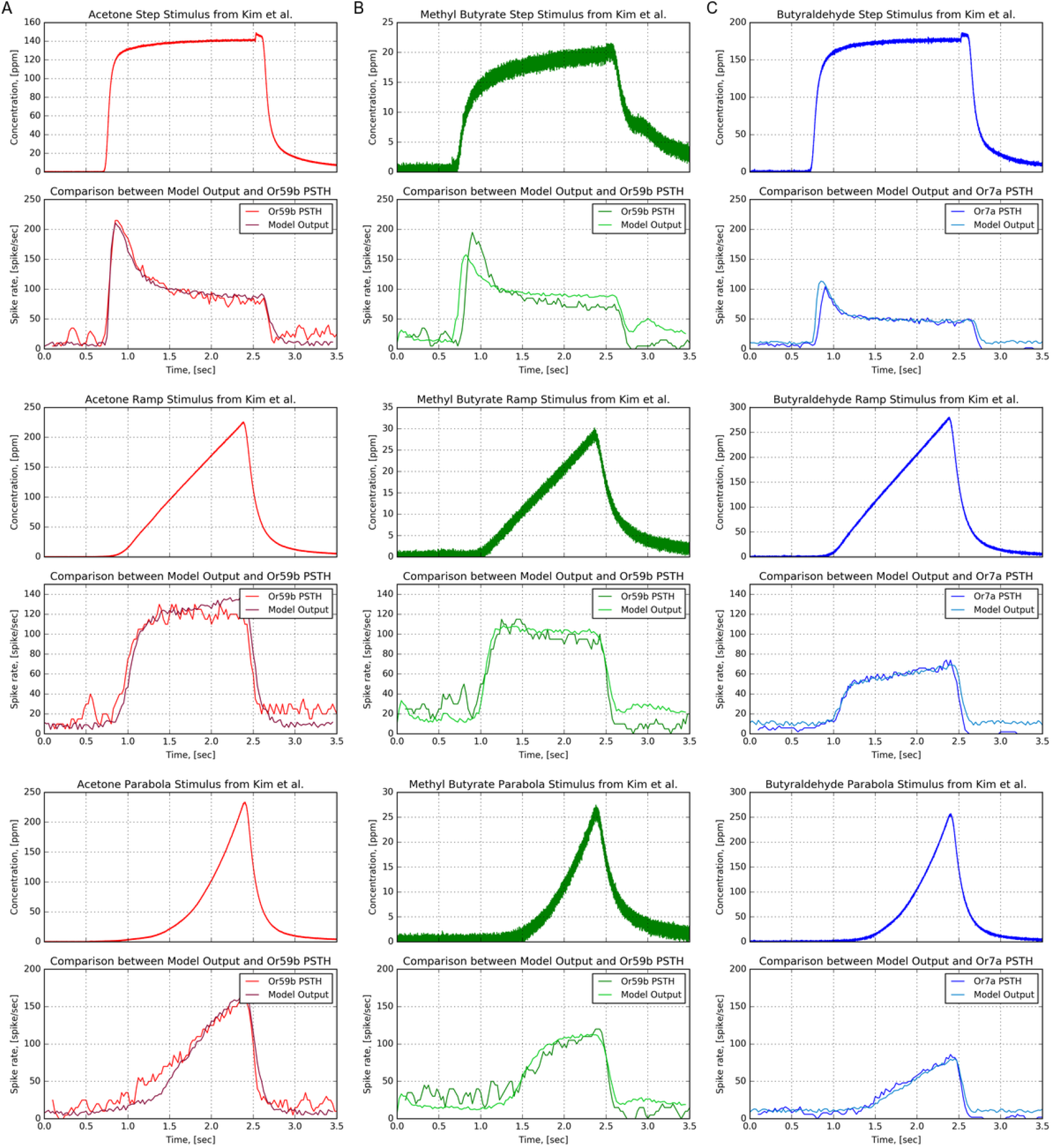
Characterization of the OTP/BSG cascade with multiple odorants and receptor types. Three odorant-receptor pairs are tested: 1) *Or59b* and *acetone*, 2) *Or59b* and *methyl butyrate*, and 3) *Or7a* OSN and *butyraldehyde*. (A) *Or59b* OSN in response to *acetone*. (B) *Or59b* OSN in response to *methyl butyrate*. (C) *Or7a* OSN in response to *butyraldehyde*. (Odd rows) Odorant stimuli. (Even rows) PSTH from the model output and experimental recordings (*Kim et al., 2011a*) (Reproduced from (*Kim et al., 2011a*) using the original raw data and the same color code.)

Here, we study the temporal response of the AMP LPU, that consists of 50 OSN groups (*Masse et al., 2009*; *Su et al., 2009*). Each of 50 OSN groups consists of 25 OTP/BSG cascades (neurons) that express an unique receptor type. We tested the AMP LPU with the same staircase waveform as in Figure 5.B. For an assumed odorant, we assigned the same odorant-receptor affinity to OTPs in the same OSN group. The value of the affinity for each of the 25 OTP ranges between 2 · 10^−4^ and 10^−2^ with a step size of 2 · 10^−4^. The dissociation rate for all OTP models was set to 10^2^. For simplicity, we used the same set of parameters for all cascades across all OSN groups. The parameters of the OTP model are given in Table 2, and the parameters of the BSG model are listed in Appendix 2. From the spike sequences generated by the 25 cascades we evaluated the PSTH for each of the 50 OSN groups.

**Figure 8.**
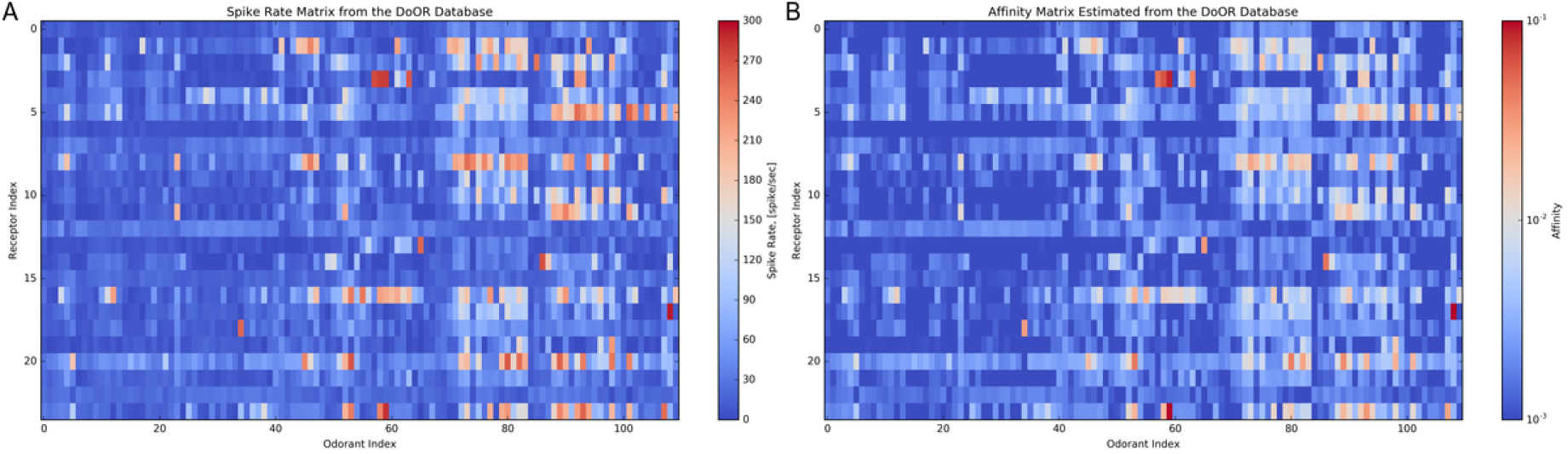
Estimating the odorant-receptor affinity matrix. (**A**) Spike rate matrix from the DoOR database containing 24 odorant receptors and 110 odorants. The data was originally published in (*Hallem and Carlson, 2006*). Each column represents an odorant, and each row represents an OSN receptor type. (**B**) Each entry of the affinity matrix is estimated from each entry of the spike rate matrix using the inverse of the function empirically determined with **Algorithm 1**. Note the log-scale color map for the affinity values.

**Figure 9.**
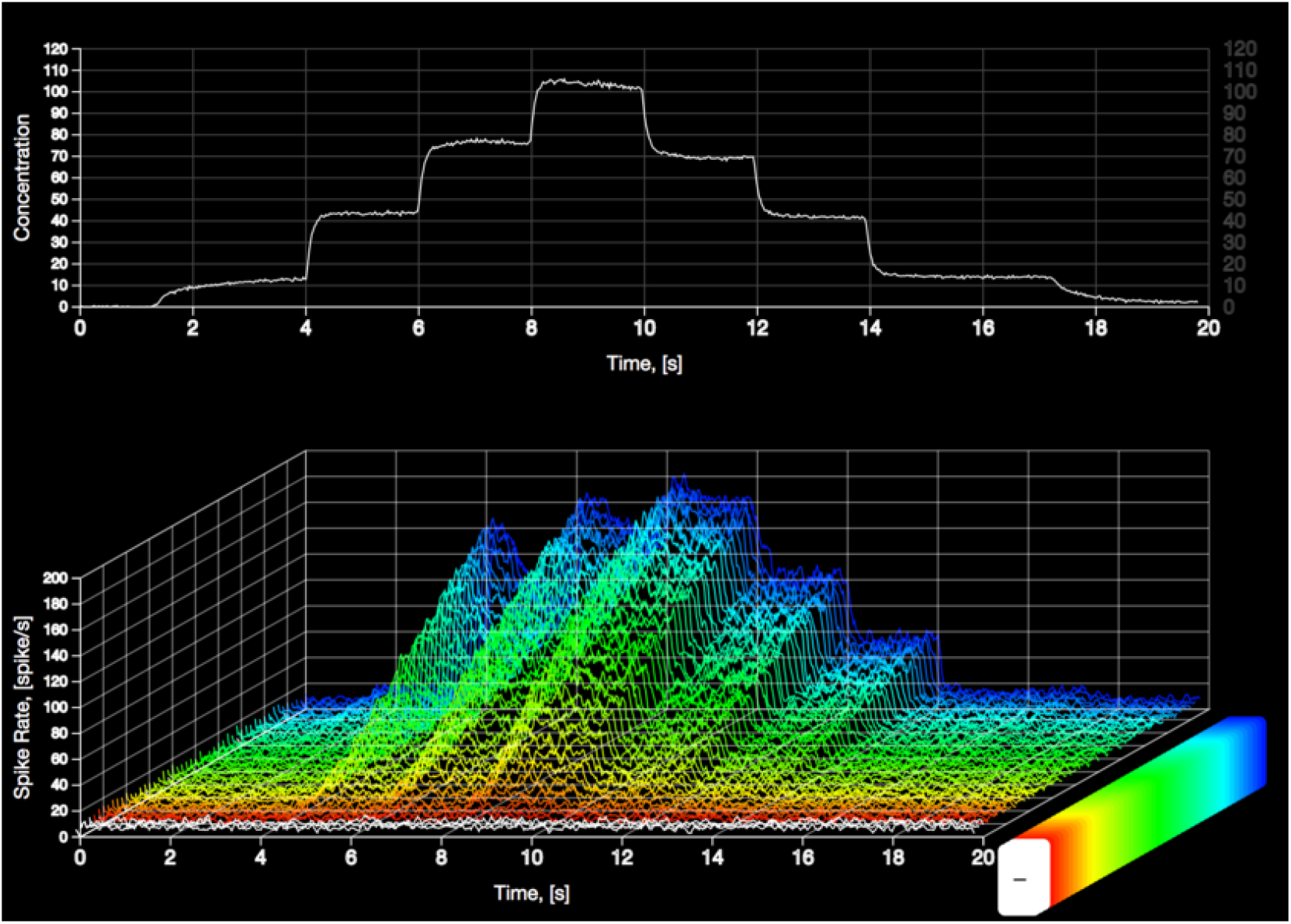
Preview of the animation demonstrating the AMP LPU in response to a staircase concentration waveform. The animation was rendered by NeuroGFX (*Ukani et al., 2019*). Each of the OSN groups consists of 25 fruit fly OTP/BSG cascades. The PSTH for each of the OSN groups was evaluated from the spike sequences generated by 25 cascades. The affinity for each of the 50 OSN groups was assumed to be ranging between 2 · 10^−4^ and 10^−2^ with a step size 2 · 10^−4^. The dissociation rate for all OTP models was set to 10^2^. The rest of parameters of both OTP and BSG are given in Table 2 and **Appendix 2**, respectively. (**top**) Staircase odorant stimulus. (**bottom**) 3D view of 50 OTP/BSG PSTHs. The response curves are sorted in ascending order according to the amplitude of the binding rate.

We visualize the 50 PSTHs and provide the preview and the link to the animation in Figure 9. The animation is rendered by NeuroGFX, a key component of FFBO (*Ukani et al., 2019*) (both NeuroGFX and FFBO are briefly presented in the *Methods and Materials* section). As shown, the top plot in the animation (and in Figure 9) shows the staircase odorant waveform, and the bottom plot shows the 3D view of the 50 PSTHs.

The resultant PSTHs exhibit distinct temporal responses across different OSN groups. Both the concentration and concentration gradient of odorants with (overall) high binding rate values is 2D encoded.

For receptors with extremely low (overall) binding rate values, the OTP/BSG cascade generates an output only after the concentration amplitude exceeds a certain value. For example, as shown in Figure 9, OSNs expressing receptors marked with orange color remain silent in the time interval between 2 to 6 seconds under a weak amplitude concentration stimulation. They start reacting to the odorant stimulus after 8 seconds as the amplitude increases from 80 ppm to 100 ppm. This closely matches the experimental recordings (*Hallem and Carlson, 2006*).

To further evaluate the AMP LPU, we used the affinity matrix estimated from the DoOR database (see Figure 8), and simulated 24 OSN groups in response to 110 different odorants. The dissociation rate for OSN groups was assumed to be 132, as it can not be estimated from the available records in the DoOR database. We applied the same staircase odorant waveform as above, and visualized the PSTH of OSN groups with an animation. In Figure 10, we provide the preview and the link to the animation. As shown, the top of the animation in Figure 10 shows the staircase odorant waveform as a function of time, and the bottom of the animation (also in Figure 10) shows the spike rate matrix for 24 OSN groups and 110 odorants at each time point. Each row of the matrix represents an OSN group, and each column of the matrix corresponds to an odorant. The animation demonstrates that the intensity of the spike rate matrix increases dramatically at each jump of the staircase waveform and drops down to a steady state value afterwards. This striking feature is due to the large value of the concentration gradient at transition times and clearly stands out in the animation.

**Figure 10.**
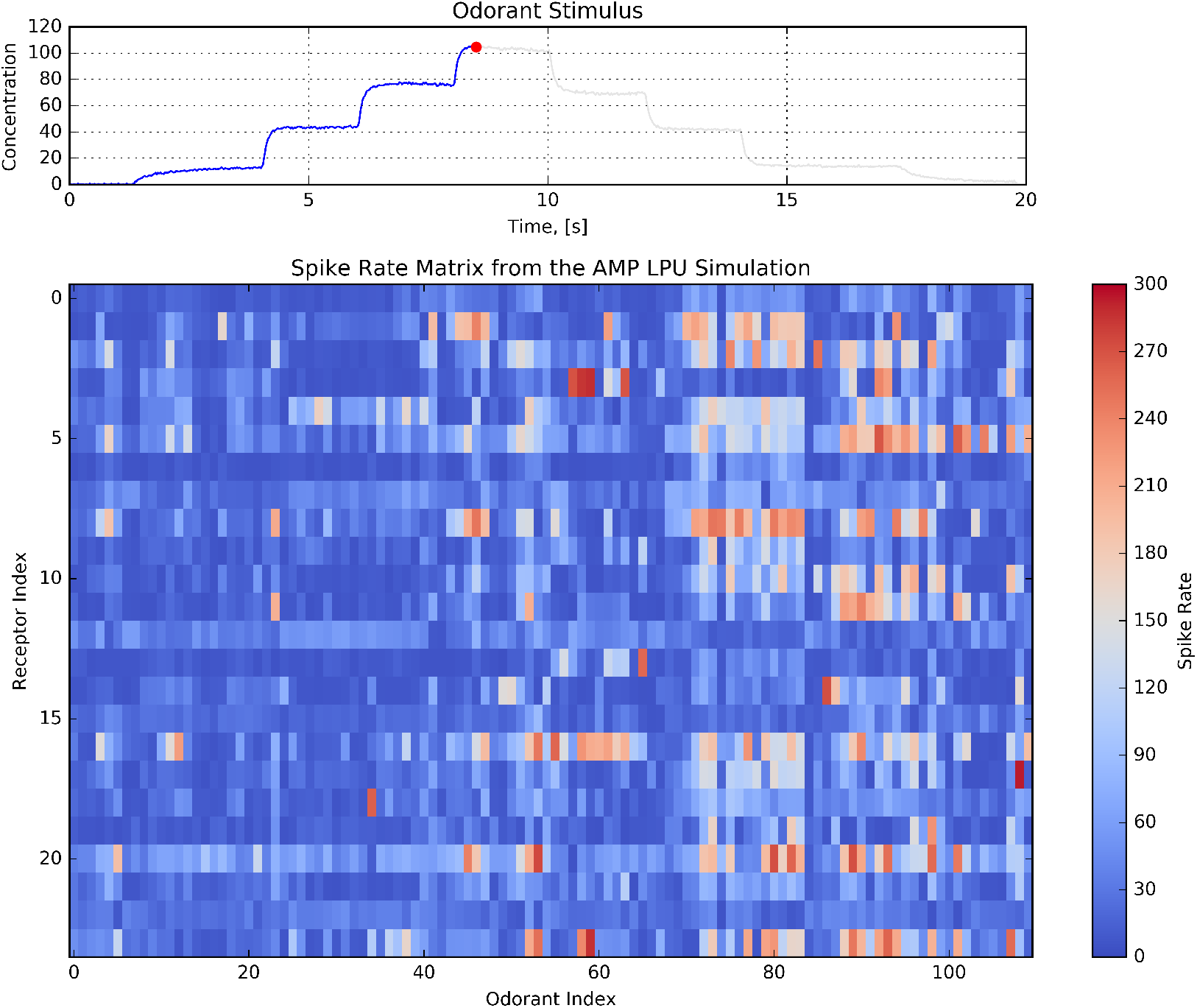
Preview of the animation of the spike rate matrix of 24 OSN groups in response to 110 odorants. Each of the OSN groups consists of 25 OTP/BSG cascades. The PSTH for each of the OSN groups is evaluated from the spike sequences generated by the 25 cascades. Each row of the matrix represents an OSN group, and each column of the matrix corresponds to an odorant. The affinity for each pairs of OSN groups and odorant is estimated using the DoOR database (see Figure 8). The dissociation of all OSN groups is assumed to be 132. (**top**) Staircase odorant waveform. (**bottom**) Dynamics of the spike rate matrix across time.

## Discussion

Successful modeling of encoding of odorants by olfactory sensory neurons spread across the antennae and maxillary palps requires means of easily constructing and testing a range of hypotheses regarding the transduction of odorants into spike trains. The essential functionality of olfactory sensory neurons that we focussed on here is their concurrent encoding of both odorant identity and odorant concentration. To address these two functional aspects we presented an in-depth description of OSNs modeled as two stage samplers that quantitatively encode both the odorant identity and its concentration profile.

We devised a class of modular OSN models as a cascade consisting of an odorant transduction process and a biophysical spike generator. The OTP model consists of an active receptor model and a co-receptor channel model. The the BSG model employed here is based on the Connor-Stevens neuron model. After developing the OTP and BSG models, our focus was on the biological validation of the OTP/BSG cascade. To validate our modeling approach, we examined the response of the fruit fly OSN model to odorant waveforms that were previously used in experiments with different odorants and receptors, and compared the model responses with electrophysiology recordings. Our results show that the OTP/BSG cascade model proposed here indeed closely matches the complex temporal response of OSNs.

### Limitation of the DoOR Database

The odorant-receptor affinity matrix together with the spike rate matrix provide the macroscopic I/O characterization of the OSN model at the population level. However, in the absence of additional information, such as the slope, width, or peak of the OSN response to the odorant onset, the dissociation rate can not be estimated with Algorithm 1 as such information is currently not available in the DoOR database. Thus, whereas previously the affinity and the dissociation rate are both estimated for multiple odorant-receptor pairs, only the affinity can be estimated for the odorant-receptor pairs recorded in the DoOR database. The dissociation rate together with the affinity are both required for reproducing the temporal response of OSNs. The latter alone can only characterize the steady state response. This illustrates some of the limitations of the DoOR datasets for characterizing the temporal response properties of OSNs, despite their richness for characterizing the steady state response of odorant-receptor pairs.

### Combinatorial Coding of Odorant Identity is Concentration Dependent

The output of simulated OSN population demonstrates that the odorant identity is encoded in the set of odorant-activated OSN groups expressing the same receptor type. Different odorants evoke different sets of OSN groups, as shown in Figure 10. More importantly, the size of the set expands or reduces as the odorant concentration increases or decreases.

### Complexity of the Input Space of Olfactory Neuropils

Our approach models on the molecular level the combinatorial complexity of the transformation taking place in *Drosophila* antennae OSNs. The transformation maps the odorant-receptor binding rate tensor modulated by the odorant concentration profile and the odorant-receptor dissociation rate tensor into OSN spike trains, respectively. The resulting *concentration-dependent combinatorial code* determines the complexity of the input space driving olfactory processing in the downstream neuropils, such as odorant recognition (*Badel et al., 2016*) and olfactory associative learning (*Lin et al., 2014*; *Hige et al., 2015*).

### Future Work

The odorant receptor and the pheromone receptor share similar temporal variability in response to input stimuli (*Gu et al., 2009*), despite the differences in protein structure and chemical signaling between the two receptor families. Therefore, the fruit fly OTP model can be extended to model pheromone receptors. Pheromones can be modeled as odorants with an extremely high single receptor binding and dissociation rates.

The active receptor model can be readily extended to odorant mixtures. One interesting question is to study the odorant encoding of OSNs in the presence of a background odorant. Another potential direction is to investigate the “cocktail party” problem of odorant mixtures (*Rokni et al., 2014*).

**Table 2.** Summary of the parameters in the fruit fly odorant transduction model.

| Variable | Value | Description |
| --- | --- | --- |
| $\alpha_1$ | $1.570 \cdot 10^1$ | cutoff frequency of the filter modeling the peri-receptor process |
| $\beta_1$ | $8.000 \cdot 10^{-1}$ | slope of the transition region of the peri-receptor process filter |
| $\gamma$ | $1.750 \cdot 10^{-1}$ | scaling factor of the filtered odorant concentration gradient |
| $\alpha_2$ | $8.877 \cdot 10^1$ | rate of activation of the gating variable of the *co-receptor* channel |
| $\beta_2$ | $9.789 \cdot 10^1$ | rate of deactivation of the gating variable of the *co-receptor* channel |
| $\alpha_3$ | $2.100 \cdot 10^0$ | rate of increase of the state variable of the calcium channel |
| $\beta_3$ | $1.200 \cdot 10^0$ | rate of decrease of the state variable of the calcium channel |
| $\kappa$ | $7.089 \cdot 10^3$ | feedback strength from the calcium channel to the *co-receptor* channel |
| $c$ | $7.534 \cdot 10^{-2}$ | value achieving the half-activation of the *co-receptor* channel |
| $p$ | 1 | the Hill coefficient of the *co-receptor* channel |
| $I_{max}$ | $7.774 \cdot 10^1$ | maximum current amplitude generated by the *co-receptor* channel |

## Methods and Materials

### I/O Characterization of the OTP/BSG Cascade

Is the OTP model given in Equation 7 capable of qualitatively reproducing transduction currents as those recorded in voltage clamp experiments? We empirically explore this question below.

We first empirically tuned the parameters of the odorant transduction process model so as to generate similar transduction currents as recorded in the voltage-clamp setup published in (*Cao et al., 2016*). The binding rates and the dissociation rates of all OTP models were set to 1 and 132, respectively, and values of the other parameters are listed in Table 2.

We evaluated the model using step stimuli *u_s_*(*t*), ramp stimuli *u_r_*(*t*), and parabola stimuli *u_p_*(*t*), chosen as,

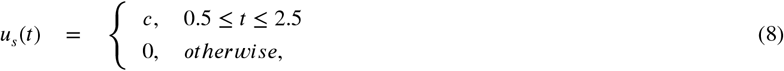

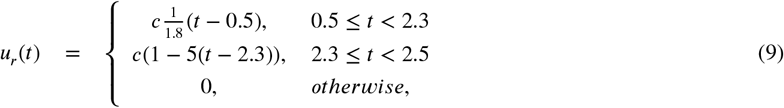

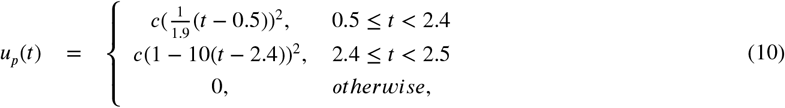

where *c* is a scalar ranging between 1 and 101 with a step size of 5.

The response at the output of the peri-receptor process *u* * *h*, the odorant concentration profile [**v**]_*ron*_, and the ratio of bound receptor [**x**_1_]_*ron*_ are shown in Figure 11. The slope of the rising phase of *u* * *h* after the onset of odorant is due to the effect of the filter *h*(*t*). The odorant concentration profile [**v**]_*ron*_ encodes the gradient of the concentration for the step stimuli (see the chair-shaped response), but less so for the ramp and parabola stimuli. Lastly, the bound receptor [**x**_1_]_*ron*_ transforms the odorant concentration profile and maps it into a bounded range between 0 and 1.

As shown in Figure 11.A, the OTP model exhibits temporal response dynamics akin to the adaption phenomena reported in (*Cao et al., 2016*). Furthermore, we tested the OTP model with ramp and parabola stimuli of different concentration amplitude values, as shown in Figure 11.

**Figure 11.**
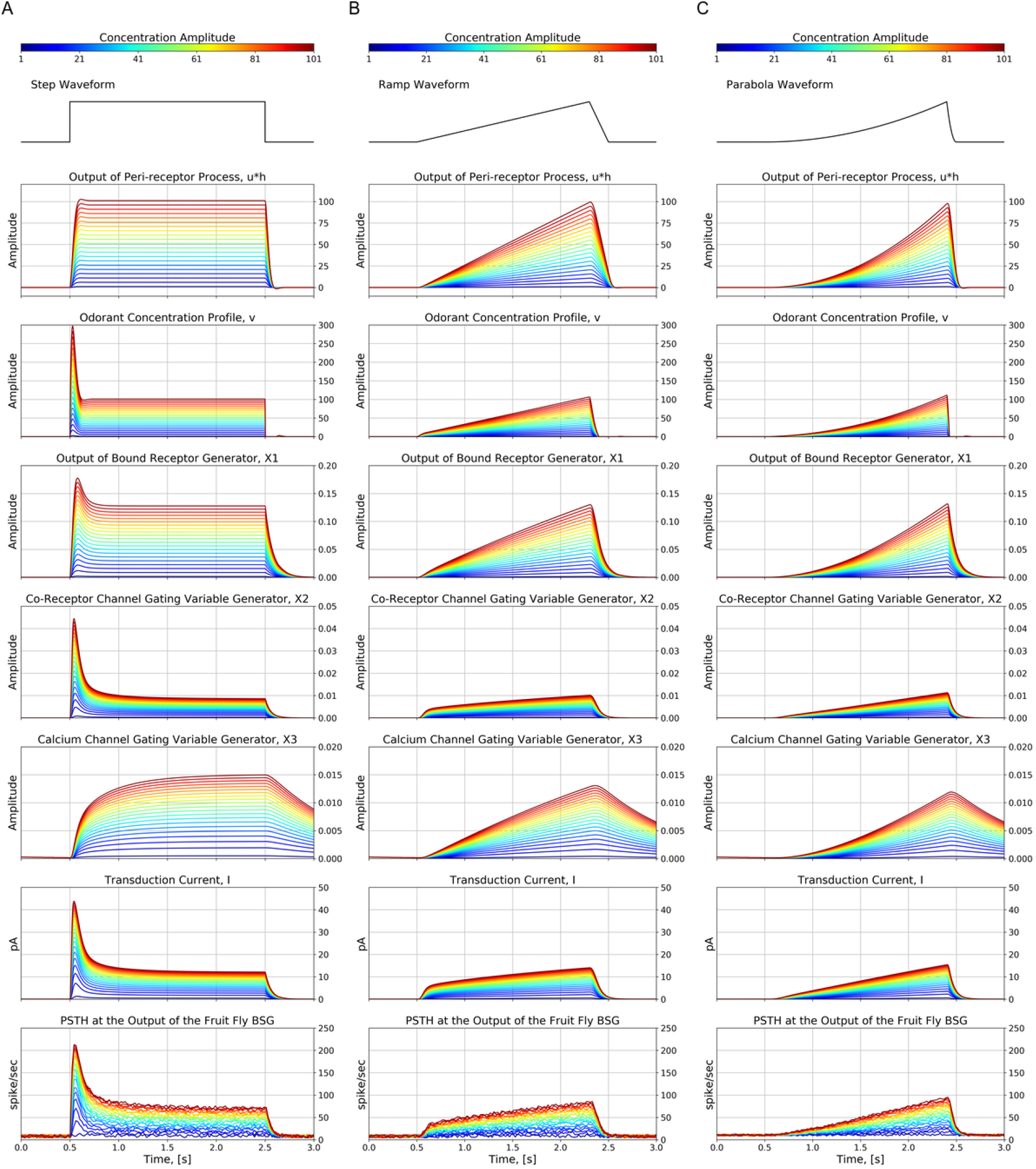
Characterization of the fruit fly OSN model in response to odorant stimuli of different concentration amplitude values, ranging between 1 and 101 ppm with a step size of 5 ppm. The parameters of the OTP model are given in Table 2, and the parameters of the BSG model are listed in **Appendix 2**. The binding rate and the dissociation rate of all OTP models were set to 1 and 132, respectively. (**A**) Step stimulus given by Equation 8. (**B**) Ramp stimulus given by Equation 9. (**C**) Parabola stimulus given by Equation 10.

Compared with the stimulus response of the active receptor model, the response of the OTP model exhibits a complex temporal variability, that is sensitive to both the amplitude and the gradient of the odorant stimulus waveform. For example, as shown in Figure 11.B, the response of the OTP model to the ramp stimulus first increases linearly as the ramp stimulus increases, but then plateaus and remains constant as the gradient of the ramp stimulus is a constant. In addition, as shown in Figure 11.C, the response of the OTP model to the parabola stimulus roughly resembles a ramp function that closely matches with the gradient of the parabola stimulus.

The complex temporal response of the OTP model is due to the feedback received by the *co-receptor channel* from the *calcium channel.* Without the *calcium channel* feedback, the OTP model is reduced to a three-stage (*peri-receptor processing, bound-receptor generator*, and *co-receptor channel*) feedforward model. The feedback enables the OTP model to encode the odorant concentration profile components, i.e., both the filtered odorant concentration and concentration gradient. In addition, the nonlinearities embedded in the current generation of the *co-receptor channel* (see also Equation 6) acts as a normalization block, that facilitates the OTP model to map a stimulus with a wide range of amplitude values into a bounded transduction current.

The temporal response variability of the OTP/BSG cascade is similar to the transduction current generated by the OTP model. The similarity between the responses of the OTP model and the OTP/BSG cascade suggests that the temporal variability of the odorant concentration profile is primarily encoded in the OTP model. The BSG model is simply a sampling device mapping input current waveforms into spike trains.

### Evaluating the Steady State Response of the OTP/BSG Cascade

We note that Equation 3 can be written as,

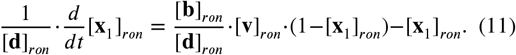

where [**b**]_*ron*_/[**b**]_*ron*_ is the odorant-receptor or ligand-receptor “affinity” (*Kastritis and Bonvin, 2013*). The active receptor model postulated in Equation 11 implies that in steady state the product between the odorant-receptor affinity and the odorant concentration profile is the main figure of merit for I/O characterization of the fruit fly OTP/BSG cascade. To study its mapping into spike rate, we simulated OTP/BSG cascades with constant stimuli, and evaluated the spike rate at steady state.

The amplitude of step stimuli ranges between 10^−1^ and 10^5^ with a step size of 0.1 on the logarithmic scale. The affinity ranges between 10^−2^ and 10^1^ with a step size of 0.01 on the logarithmic scale. The parameters of the OTP model are given in Table 2, and the parameters of the BSG model are listed in **Appendix 2**. The step stimulus is 5 second long, and the OTP/BSG cascades reach steady state roughly after 3 seconds. We calculated the spike rate using a window between 4 to 5 seconds, and plotted the results in 2D in Figure 12. Note that the *x*-axis in Figure 12 is on the logarithmic scale. As shown in Figure 12, for different values of the odorant-receptor affinity, the mapping of the concentration amplitude into spike rate shifts along the x-axis. A low affinity value requires a higher concentration amplitude value in order to elicit spikes above the spontaneous activity rate.

**Figure 12.**
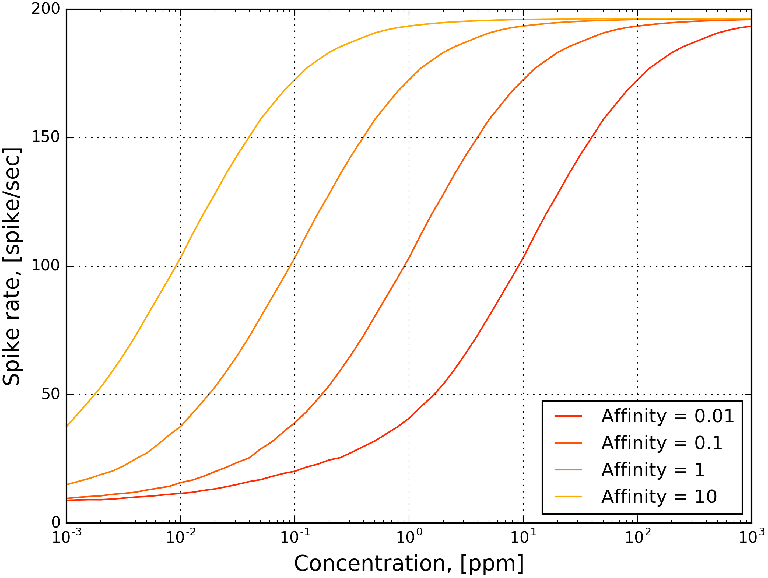
The transformation of the odorant concentration amplitude into steady-state spike rate by the OTP/BSG cascade for fixed values of the ligand-receptor affinity in response to 5-second-long constant stimuli. The parameters of the OTP model are given in Table 2, and the parameters of the BSG model are listed in **Appendix 2**. The amplitude of the constant odorant stimuli ranges between 10^−3^ and 10^3^ with a step size of 0.1 on the logarithmic scale. The spike rate is calculated in a window between 4 and 5 seconds.

As shown in Figure 12, the transformation of the product between the odorant-receptor affinity and the concentration amplitude into spike rate resembles a sigmoidal function. The OTP/BSG cascade starts spiking only after the product exceeds a certain threshold value. For odorant-receptor pairs with a low affinity, the firing activity requires a larger minimal amplitude of concentration than for those with a higher affinity value. This, again, coincides with experimental findings that odorant-receptor pairs with lower affinity require higher odorant concentration values in order to elicit spiking activity (*Hallem and Carlson, 2006*).

### Reproducing the 2D Encoding of the OSNs

To examine whether the fruit fly OTP/BSG cascade exhibits the 2D encoding property, we stimulated the cascade with the set of 110 triangular concentration waveforms that were previously used in experiments (*Kim et al., 2011b*) with *Or59b* and *acetone.* The triangular waveforms and their trajectories are plotted in Figure 13.A and Figure 13.C. We applied each of the triangular waveforms to 25 OTP/BSG cascades, and evaluated the PSTH using the spike train of all 25 cascades with a 20 ms bin size and 10 ms time shift between consecutive bins. The binding and dissociation rates of all OTP/BSG cascades was set to 3.141 · 10^4^ and 1.205 · 10^1^, respectively. The parameters of the OTP model are listed in Table 2, and the parameters of the BSG model are listed in **Appendix 2**.

The responses of the OTP/BSG cascade are given in Figure 13. The PSTH of the OTP/BSG cascade in response to different waveforms is color-coded in both the 2D and 3D view, as shown in Figure 13.B and Figure 13.D, respectively. In addition, we applied the 2D ridge regression algorithm to identify a 2D encoding manifold that best fits the PSTHs. The manifold and its contour are depicted in Figure 13.F and Figure 13.E, respectively. Similarly to the case of the staircase waveform, the OTP/BSG cascade firing rate increases dramatically as the concentration increases.

As shown in Figure 13.F, a 2D encoding manifold in a concentration and concentration gradient space provides a quantitative description of the OTP/BSG cascade. Examining Figure 13.F, we note that the 2D encoding manifold is highly nonlinear and that the OTP/BSG cascade clearly encodes the odorant concentration and its rate of change. The OTP/BSG cascade responds very strongly to even the smallest positive values of the gradient and encodes only positive concentration gradients at low odorant concentrations. At high concentrations the OSN mostly encodes the odorant concentration.

**Figure 13.**
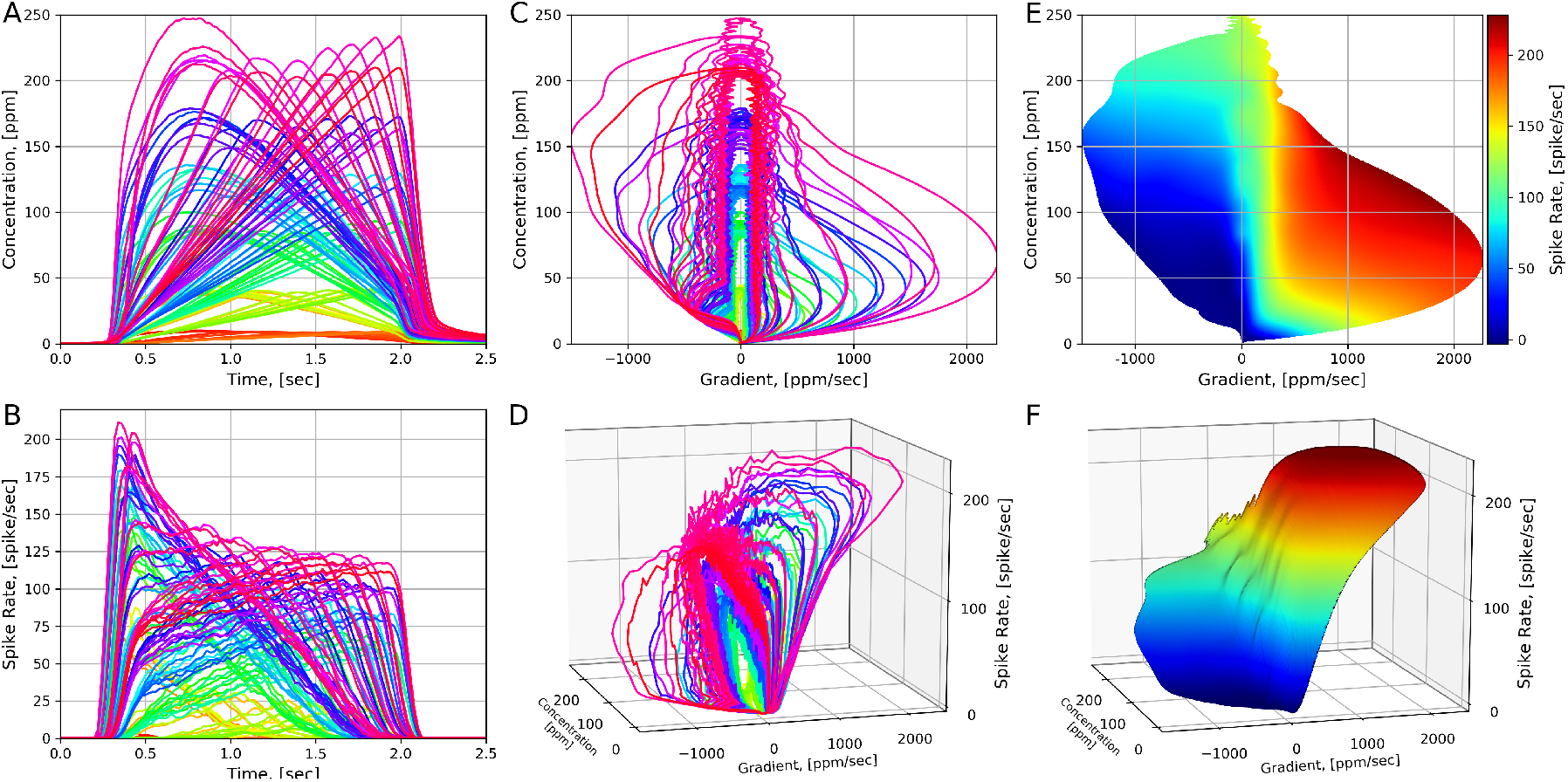
Characterizing the 2D encoding of the OTP/BSG cascade. (**A**) 110 triangular concentration waveforms. Different colors correspond to distinct triangular waveforms. (**B**) The PSTHs of the OTP/BSG cascade in response to triangular concentration waveforms. Different colors correspond to distinct waveforms. PSTHs were computed using a 20 ms bin size and a 10 ms time shift between consecutive bins. (**C**) The trajectories of triangular waveforms plotted in the concentration and concentration gradient plane. (**D**) The trajectories of PSTHs plotted in the concentration and concentration gradient plane. (**E**) The contour plot of the 2D manifold. (**F**) The 2D Encoding Manifold fitted to the trajectories of PSTHs. The manifold is generated by applying a 2D ridge estimator to the PSTHs.

### Estimation Algorithm for Affinity Value and Dissociation Rate

The receptor expressed by an OSN encodes an odorant as the pair ([**b**]_*ron*_ · [**v**]_*ron*_, [**d**]_*ron*_), i.e., the product of the odorant-receptor binding rate and the odorant concentration profile, and the odorant-receptor dissociation rate. The OTP/BSG cascade then samples and presents this representation as a train of spikes. As shown in Equation 11, for a constant stimulus with amplitude *u*, the pair is equivalent to 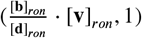. In addition, we also note that [**d**]_*ron*_ · *dt* = *d*([**d**]_*ron*_*t*) is, in effect, a time change.

An algorithm to estimate the values of the odorant-receptor biding and dissociation rates may, therefore,

- estimate the ligand-receptor affinity in steady state when the LHS of Equation 11 is zero for all values of the dissociation rate [**d**]_*ron*_, and
- estimate of the dissociation rate [**d**]_*ron*_ during a concentration jump assuming the value of the ligand-receptor affinity to be the one obtained in 1. above.

We describe the procedure above in more detail in Algorithm 1.

#### Algorithm 1 Estimation of the Affinity, Binding and Dissociation Rates

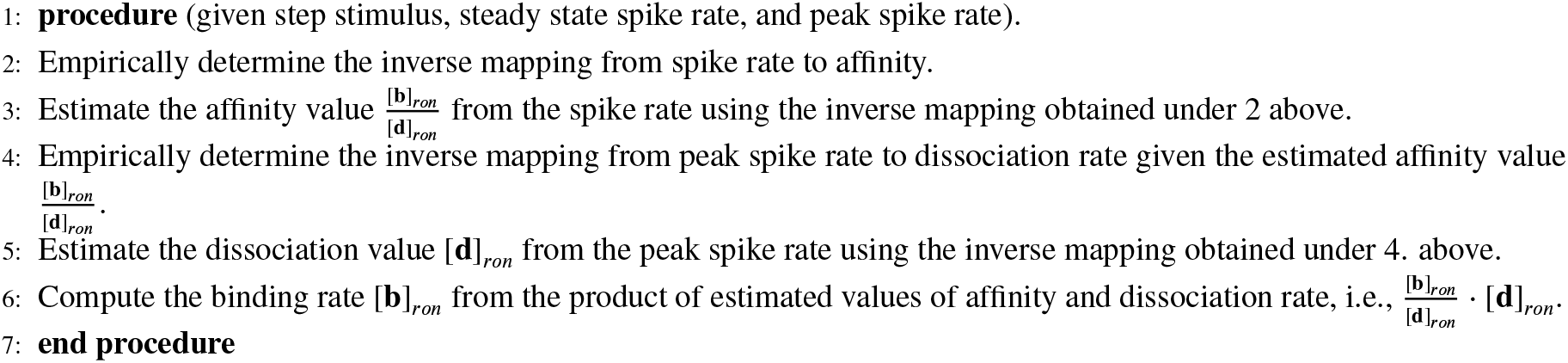

### Numerical Stability of the OSN Model

It is easy to see that [**x**_1_]_*ron*_ and [**x**_2_]_*ron*_ take values in [0,1]. This is because the value of the derivative 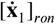 at [**x**_2_]_*ron*_ = 0 is positive and the derivative 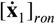 at [**x**_1_]_*ron*_ = 1 is negative. Same reasoning applies to 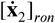. Finally, we also note that [**x**_3_]_*ron*_ is positive.

### Simulation Setup in the Fruit Fly Brain Observatory Platform

The Fruit Fly Brain Observatory (FFBO) (*Ukani et al., 2019*) is an open-source platform for the emulation and biological validation of fruit fly brain models in health and disease. It provides users highly intuitive tools to execute neural circuit models on modern commodity hardware (*Givon and Lazar, 2016*). NeuroGFX is the key FFBO component supporting the implementation of AMP LPU. NeuroGFX conjoins the simultaneous operations of model exploration and model execution with a unified graphical web interface that renders interactive simulation results.

We evaluated the response to each of the stimuli by 50 neuron groups, each group consisting of the same OSN type. Each of 50 groups consisted of 25 OSNs, and in total there were 1, 250 OSNs. We then computed the PSTH for each of OSN groups using the resultant 25 spike sequences in each of the groups. The PSTH had a 20 ms bin size and was shifted by a 10 ms time interval between consecutive bins. The parameters of all OTP models are given in Table 2. The binding rate was separately set for each odorant-receptor pair. We used the same set of parameters for all 1250 cascades, but generated different sample paths for the Brownian motion term W in Equation 12. The parameters of the BSG model are listed in Appendix 2.

## Acknowledgments

The research reported here was supported in part by AFOSR under grant #FA9550-12-1-0232 and in part by NSF under grant #1544383. We thank Anmo J. Kim for his assistance in sorting out some of the unpublished olfactory sensory neuron recordings presented here for the first time and that are part of a data trove not reported in (*Kim et al., 2011a*), (*Kim et al., 2011b*) and (*Kim et al., 2015*).

## Appendix 1 Peri-receptor Processing Model

*h*(*t*) in Equation 2 is the impulse response of a low-pass linear filter that is usually defined in the literature in frequency domain. Alternatively, *h*(*t*) is the solution to the second-order differential equation,

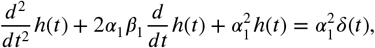

with the initial condition *h*(0) = 0 and *dh/dt*|_*t*=0_ = 0, where *δ* is the Dirac-function. The value of *α_1_* and *β*_1_ are given in Table 2, and the corresponding *h*(*t*) has an effective bandwidth of 15 Hz.

## Appendix 2 Biophysical Spike Generator Model

We restrict our choice of the spiking mechanism of OSNs to biophysical spike generators (BSG) such as the Hodgkin-Huxley, the Morris-Lecar, and the Connor-Stevens point neuron models. For simplicity of presentation, we only describe here the Connor-Stevens (CS) neuron model (*Connor and Stevens, 1971*). The CS model can be expressed in compact form as

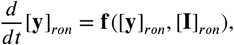

with **y** = [*V, n, m, h, p, q*]^*τ*^ is a vector of state variables, **f** is a vector function of the same dimension, and **I** is the transduction current generated by the OTP model. Here *ron* takes the same values as the same subscript in the OTP model. Compared with the classic Hodgkin-Huxley neuron model, the CS neuron model has a continuous F-I curve (*Gabbiani and Cox, 2010*), and is capable of encoding weak pulse stimuli with low spiking rates. It also has a wide spiking rate range that sufficiently covers the spiking rate range of the OSNs.

The CS neuron model does not fire spontaneously, and requires a minimum value of the input current to trigger firing. OSNs are noisy and fire spontaneously on average 8 spikes/s. To mitigate this mismatch, we added noise to the CS neuron model,

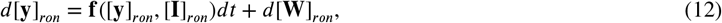

where 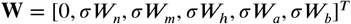, and (*W_n_, W_n_, W_h_, W_p_, W_q_*) are zero mean, unit variance independent Brownian motion processes, and *σ* is a scalar. We empirically determined the value of *σ* to be 2.05 by sweeping its value in the range of (0, 2.5) so that the noisy CS model fires some 8 spikes per second. The F-I curve of the CS neuron model for different values of *σ* is shown in Figure 1.

The CS model is based on the classic Hodgkin-Huxley (HH) neuron model (*Hodgkin and Huxley, 1952*) and can be expressed in compact form as

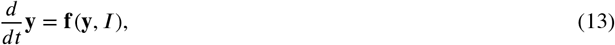

where **y** = [*V, n, m, h,p, q*]^*τ*^ is a vector of state variables, **f** is a vector function of the same dimension. For simplicity, Equation 13 omits the subscript notation in Equation 12. Similarly to the HH neuron model, the state variable *n* is a gating variable representing the activation of the potassium channel, while the state variables *m* and *h* are gating variables representing the activation and deactivation of the sodium channel, respectively. Furthermore, the variables *p* and *q* are the gating variables representing the activation and deactivation of the “a”-channel. In more detail, Equation 13 is given by

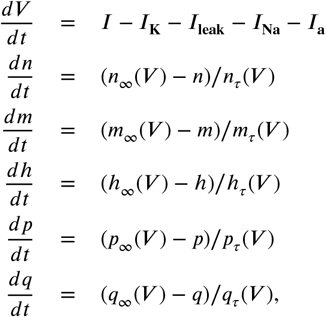

where *V* denotes the membrane voltage of the neuron model, and *I* denotes the external current. Finally,

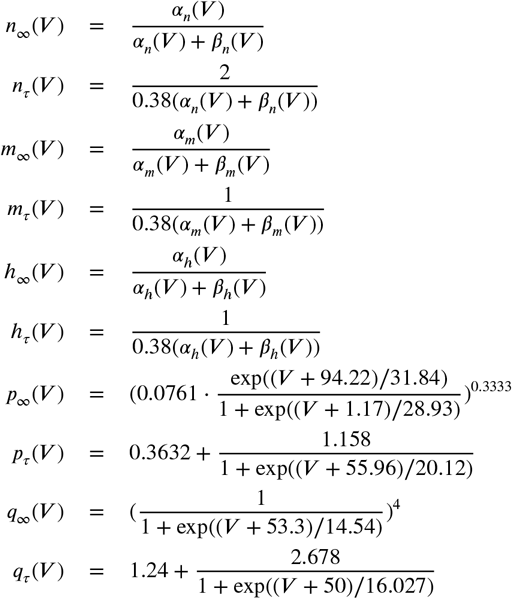

and,

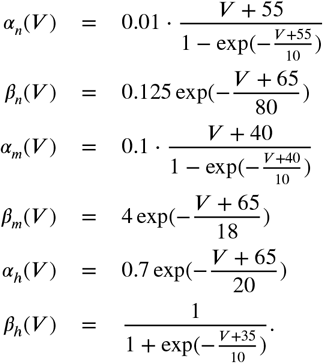

Lastly, the current generated by each of the channels is given by,

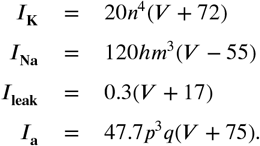

**Appendix 2 Figure 1.**
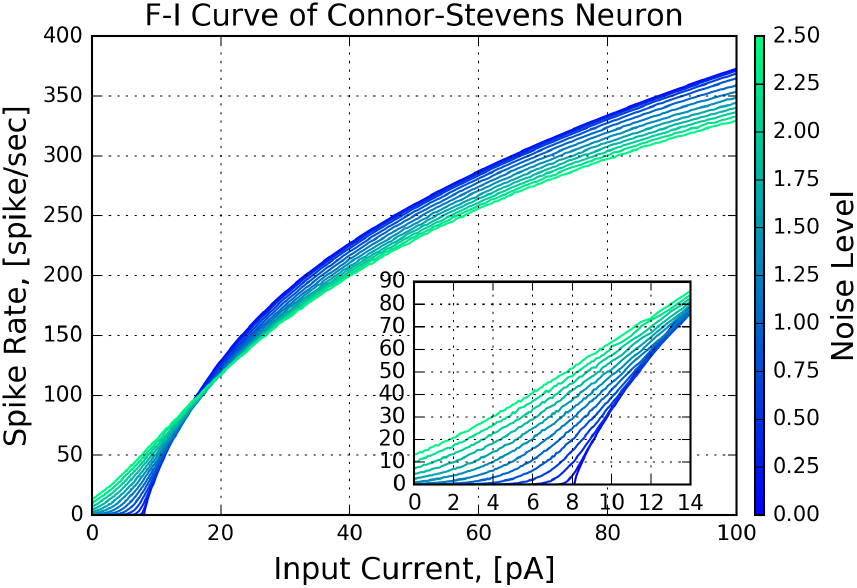
Characterization of the Connor-Stevens neuron model. The F-I curves of the model are color-coded for different noise levels (σ in Equation 12).

## Appendix 3 Two (*Acetone, Or59b*) Datasets

**Appendix 3 Figure 1.**
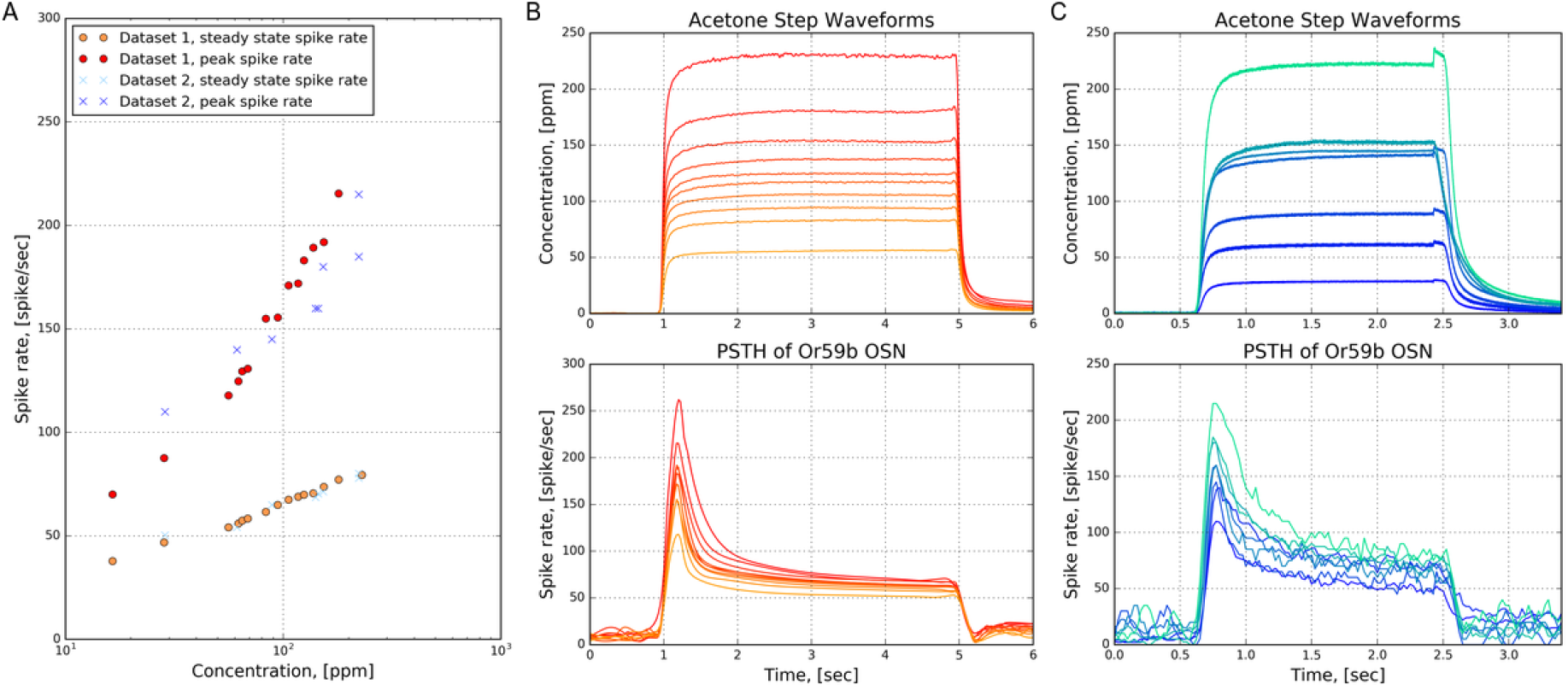
Two datasets of PSTHs of *Or59b* in response to acetone step waveforms. (**A**) The peak and steady state spike rate as a function of concentration amplitude. (**B**) Dataset 1. (top) acetone waveforms. (bottom) PSTHs of *Or59b* OSNs. (**C**) Dataset 2. (top) acetone waveforms. (bottom) PSTHs of *Or59b* OSNs.

Each of the two datasets contains the PSTHs obtained from the response of OSNs expressing *Or59b* to acetone step waveforms with different concentration amplitudes. The peak and steady state spike rate as a function of concentration amplitude for both datasets are given in Figure 1.A. The acetone step waveforms and the corresponding PSTHs of *Or59b* OSN for the two datasets are shown in Figure 1.B-C.

The two datasets are part of a repository of electrophysiology recording data for the olfactory system of the fruit fly. The details of the electrophysiology recordings setup and the odorant delivery system are given in (*Kim et al., 2011b*) and (*Kim et al., 2015*). The first dataset is made public here, while the second dataset was previously published in (*Kim et al., 2015*). The PSTH of the first dataset was computed using a 100 ms bin size and shifted by 25 ms between consecutive bins.

